# Cortical representations of languages during natural dialogue

**DOI:** 10.1101/2023.08.21.553821

**Authors:** Masahiro Yamashita, Rieko Kubo, Shinji Nishimoto

## Abstract

Individuals integrate their own words, their partner’s words, and the context of dialogue into coherent meanings. Research suggests that mutual understanding between speakers and listeners is supported by a shared representation between language production and comprehension^1,2^. However, it remains unknown how languages are represented in the brain during dialogue, which is characterized by dynamic, adaptive behaviours such as turn-taking^3,4^. Here, we used functional magnetic resonance imaging (fMRI) to compare language production and comprehension maps obtained from natural dialogue in the same participants to show that separate representations exist for language production and comprehension. While production showed selectivity towards the motor system and comprehension towards the auditory system, both production and comprehension were represented in broadly distributed regions. These separate representations were similar in amodal brain regions that integrate semantic^5,6^ and pragmatic information^7,8^, and provide a common ground for mutual understanding^1,2^, reflecting dynamic, complementary roles in interactive language use, including turn-taking^3,4,9^, backchannels^10^, and fillers^11^. Our findings suggest that separate and analogous linguistic representations for production and comprehension are interwoven in the same networks that underlie complementary interactions and making sense in dialogue.

---

Humans can construct meaning through dialogue, integrating their own words and those of their partners in a particular situation. For successful dialogue, the interlocutors must share cultural, physical, and linguistic knowledge for mutual understanding^12^, suggesting a shared neural representation of linguistic information between them^13^. The interactive alignment account^13^ hypothesizes that interlocutors automatically align linguistic representations at multiple levels, from phonological, syntactic, semantic, and situation models^14^. Neural coupling has been consistently observed between a narrative speaker and a listener in amodal brain regions^1,2^ that are involved in semantic^5,6^ and pragmatic representations^7,8^. These studies suggest that shared linguistic representations between individuals support their mutual understanding in monologue. In contrast, within individuals, linguistic representations for production and comprehension during natural dialogue remain elusive. Human dialogue is characterized by complementary roles between interlocutors, such as taking turns and planning their speech during comprehension^4,9,15^.

Furthermore, to achieve collective intelligence, people must distinguish their own perspective from that of others^16,17^. Given these human language competencies, we hypothesize that language production and comprehension are represented separately in the brain, while maintaining some degree of similarity for sense-making. In the present study, we tested this hypothesis by directly comparing linguistic representations estimated from a natural dialogue experiment using the same participants.

Studying the linguistic representations of natural dialogue in the brain is challenging because of difficulties with experimental controls and the vast vocabulary of natural language^18–20^. However, recent advances in large language models^21,22^ and voxel-wise encoding models^23,24^ offer a framework to understand how linguistic information is represented in the brain^5,6,25–27^. We used this approach to model linguistic representations of unscripted dialogue using fMRI and compared the representations for production and comprehension. To achieve this, we collected over two hours of dialogue from eight participants and modelled the linguistic representations using whole-brain blood-oxygen-level-dependent (BOLD) responses (**Fig. 1a**). The estimated representations were tested for commonality and similarity within individuals and then mapped onto the cerebral cortex using a low-dimensional space projection.

**Fig. 1.**
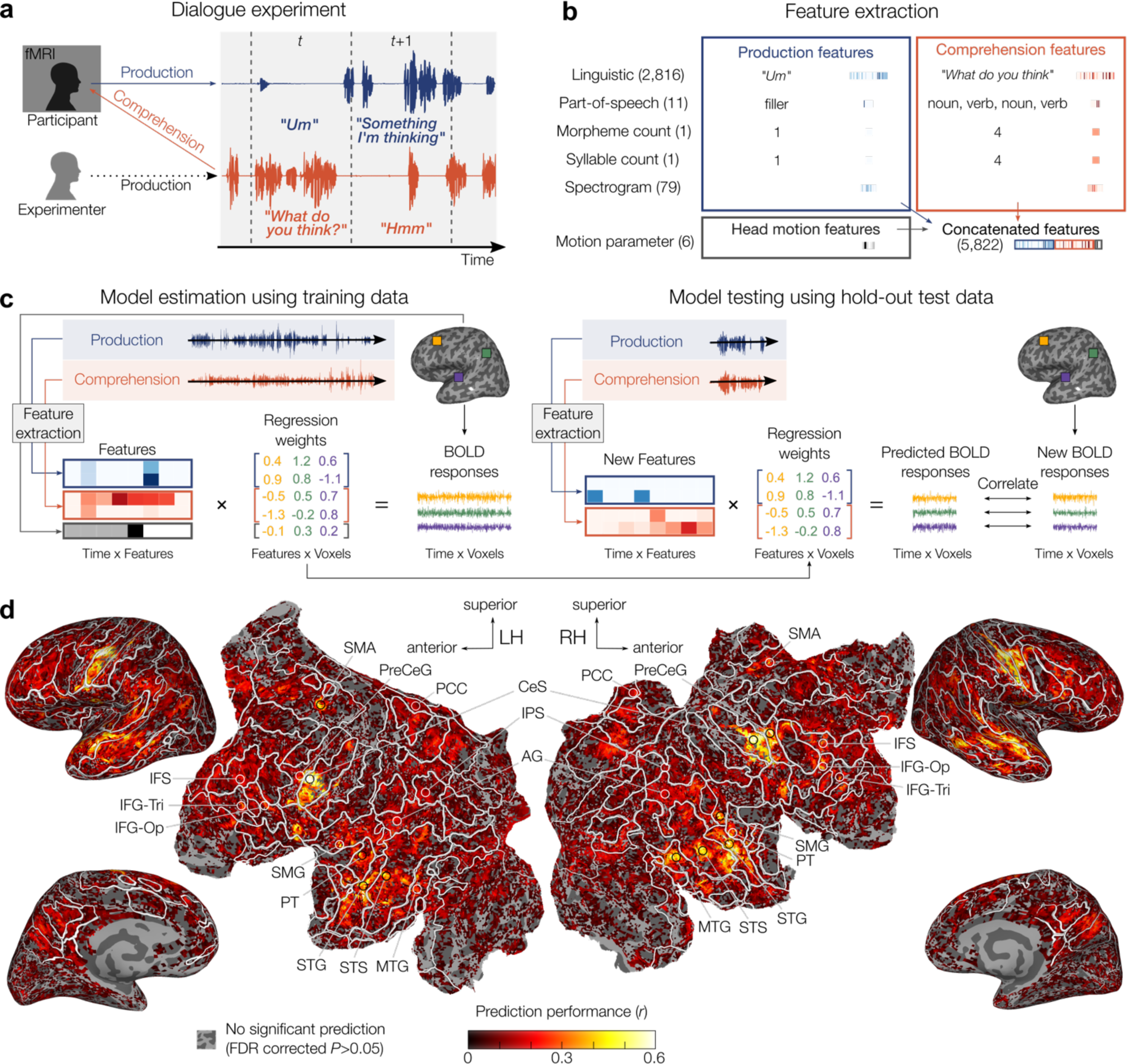
Voxel-wise model estimation and testing. **a**, Each participant engaged in a topic-restricted dialogue with the experimenter, during which brain activity was measured with fMRI. Features for production and comprehension were extracted separately from the transcribed speech content within each fMRI volume. **b**, The features include linguistic (GPT-Neox, 2,816), part-of-speech (11), morpheme count (1), syllable count (1), spectrogram (79), and head motion (6). All these features from both production and comprehension were concatenated into a voxel-wise encoding model (5,822). **c**, Using the concatenated features, the BOLD responses of each voxel were predicted separately by ridge regression. The data collected in an independent session were used to predict brain activity, and the correlation coefficient was used as the prediction accuracy (leave-session-out cross validation). **d**, Prediction performance in one participant (P8). High prediction accuracy was observed in the motor and auditory systems, with other significant predictions observed in the amodal frontal, temporal, and parietal cortices. All the participants showed similar results (**Extended Data Fig. 1**). LH: left hemisphere, RH:

## Voxel-wise model estimation and testing

In voxel-wise modelling, voxel-based encoding models predict BOLD responses in each voxel evoked by features of interest^23,24^. To extract contextualized linguistic embeddings for the utterances in the dialogues, we used a pre-trained conversational large language model, GPT-Neox (https://huggingface.co/rinna/japanese-gpt-neox-3.6b-instruction-sft). This language model was pre-trained to predict the next word from the previous context^21^ and then fine-tuned to serve as an instruction-following conversational agent. Recent fMRI studies have confirmed that GPT-based models similar to this one can elucidate brain activity evoked by linguistic content^6,27,28^. We extracted a 2,816-dimensional numerical vector from each utterance for each fMRI volume (TR = 1,000 ms, **Fig. 1a**). To account for low-level linguistic properties such as part-of-speech and spectrogram, we included additional features in the encoding model estimation and then discarded them for model testing (**Fig. 1b**). Finally, we combined the linguistic (GPT-Neox) and other low-level features from both production and comprehension (**Fig. 1b**).

To estimate how linguistic features affect BOLD responses in each cortical voxel for each participant, we used regularized linear regression. We then tested the model by applying the estimated regression weights to a new stimulus matrix not used in the model estimation (**Fig. 1c**). Here, the other low-level features were removed, and only the linguistic features were used for prediction. We found accurate predictions (*P* < 0.05, false discovery rate (FDR) corrected; **Fig. 1d** and **Extended Data Fig. 1**) in amodal frontal, temporal, and parietal regions including the superior temporal sulcus (STS), middle temporal gyrus (MTG), angular gyrus (AG), supramarginal gyrus (SMG), inferior frontal gyrus/sulcus (IFG/IFS), temporo-parietal junction (TPJ), medial prefrontal cortex (MPFC), posterior cingulate cortex (PCC), middle frontal gyrus (MFG), and intraparietal sulcus (IPS). These brain regions have been associated with semantic and pragmatic representations^5–8,25,26,29^, with shared representations between production and comprehension^1,2^, and with domain-general cognitively demanding tasks^30^. Furthermore, we found accurate predictions in sensorimotor, auditory, and visual areas, including the central sulcus (CeS), supplementary motor area (SMA) within the superior frontal gyrus (SFG), pre/postcentral sulcus/gyrus (PreCeS/PreCeG/PostCeS/PostCeG), superior temporal gyrus (STG), and planum temporale (PT). These areas have been implicated in speech processing and representation particularly in low-level phonological and articulatory processes^2,25,31–33^. The findings suggest that linguistic information for production and comprehension is represented in widely distributed regions.

## Cross-modal prediction

Next, we tested the hypothesis that linguistic representations are separate for production and comprehension within individuals in natural dialogue. To do this, we tested the generalizability of the estimated model after exchanging the weights between production and comprehension. Specifically, we exchanged the estimated weights between production and comprehension, and then examined the prediction performance (**Fig. 2a**). Again, the other low-level features (e.g., part-of-speech) were removed from the prediction. If the linguistic representations were separate, the prediction performance would be worse. Consistent with the hypothesis, we found that only a small number of voxels showed significant predictions (*P* < 0.05, FDR corrected; **Fig. 2b** and **Extended Data Fig. 2**). Comparing the prediction performance in voxels that showed significant prediction within the same modality (**Fig. 1d**), the cross-modal prediction performance significantly declined (**Fig. 2c**). These results suggest that linguistic information for production and comprehension is distinctly represented within an individual brain.

**Fig. 2.**
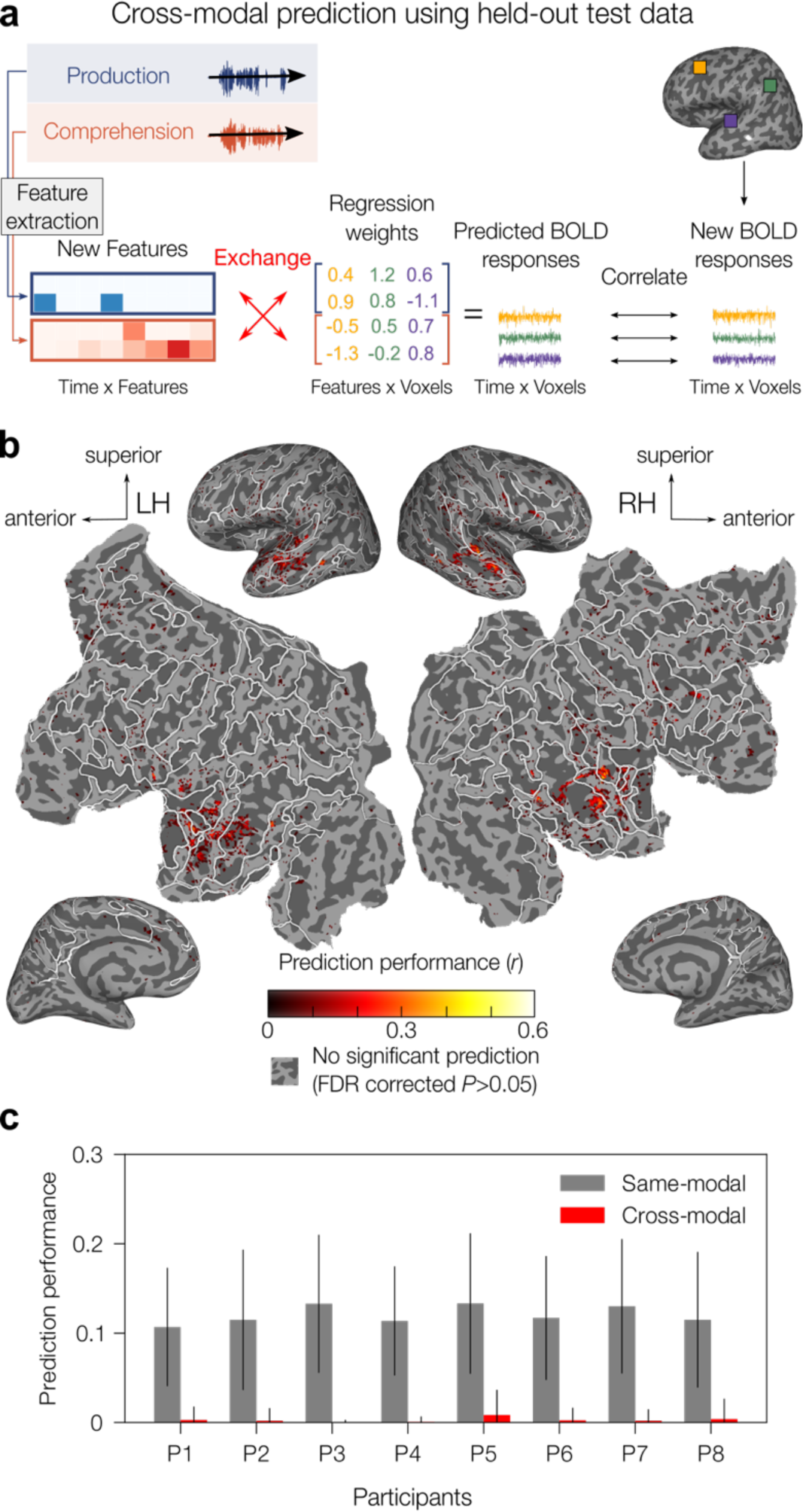
Cross-modal prediction. **a**, To test the commonality of linguistic representations between production and comprehension, we performed cross-modal prediction by combining already learned weights with stimuli of another modality (i.e., production weights and comprehension features and vice versa). **b**, Prediction performance in a participant (P8). Only a few voxels (3.9%) showed significant correlations in this participant. For this participant, voxels in bilateral superior temporal regions made significant predictions. All the other participants also showed low prediction performance (0.85-11.93% of voxels, **Extended Data Fig. 2**). **c**, Bar plot of prediction performance in the same-modal prediction and cross-modal prediction for each participant. Here, significantly predicted voxels in the same-modal prediction are included in the analysis. Error bar indicates standard deviation across the significantly predicted voxels in

## Mapping linguistic selectivity by voxel

Previous studies have shown that spatially overlapping brain regions process both language production and comprehension^1,2,34,35^, suggesting shared representations between language production and comprehension in the same networks. Next, we addressed the question of how much linguistic information for both production and comprehension is represented in the same networks within individual brains. To do this, we directly compared the estimated encoding models to investigate which modality (i.e., production or comprehension) was selectively represented for each voxel. We used two methods to map linguistic selectivity. First, we used unique variance partitioning^25,27^ to map how much variance each feature uniquely explained. Relative complement was calculated as the difference between the total variance explained by the full model and another model that removed a feature (e.g., part-of-speech for comprehension). We found that all the low-level features had only a negligible effect on the prediction performance (**Extended Data Fig. 3**). To reduce the model size, we built a reduced encoding model that concatenated linguistic (GPT-Neox) features for production and comprehension (5,632 dimension). Additionally, for the nested models, we built encoding models that use single linguistic features each for production and comprehension (2,816 dimensions). Unique variance explained by (i) production, (ii) comprehension, or (iii) intersection of production and comprehension was calculated using these models. Unique variance explained by linguistic production or comprehension was plotted onto a participant’s cortical map (**Fig. 3a**). We found production selective voxels in the frontal and parietal cortex, including bilateral CeS and IPS (shown in blue in **Fig. 3a** and **Extended Data Fig. 4**), and comprehension selective voxels in the prefrontal and temporal cortex, including the IFS and STG/STS (shown in red in **Fig. 3a** and **Extended Data Fig. 4**). These results support the classical framework of the separation of production and comprehension^36^. Second, to gain further insight into this observation, we determined the best variance partition for each significantly predicted voxel. To do this, we classified voxels into three types: variance was best explained by (i) production only, (ii) comprehension only, or (iii) production-comprehension intersection (bimodal). The intersection explained the most variance in distributed brain regions including the CeS (shown in green in **Fig. 3b** and **Extended Data Fig. 3**). We also found production explained most variance in the fronto-parietal network (FPN, shown in blue), while comprehension explained most variance in the auditory and prefrontal cortex (shown in red). Counting the number of voxels for each type in each participant, we found that the number of bimodal selective voxels was greater than the number of unimodal selective voxels in all but two participants (**Fig. 3c** and **Extended Data Fig. 3**). These results suggest that many brain regions process linguistic information for both production and comprehension in natural dialogue, providing unique evidence that these linguistic processes are interwoven in the same networks.

**Fig. 3.**
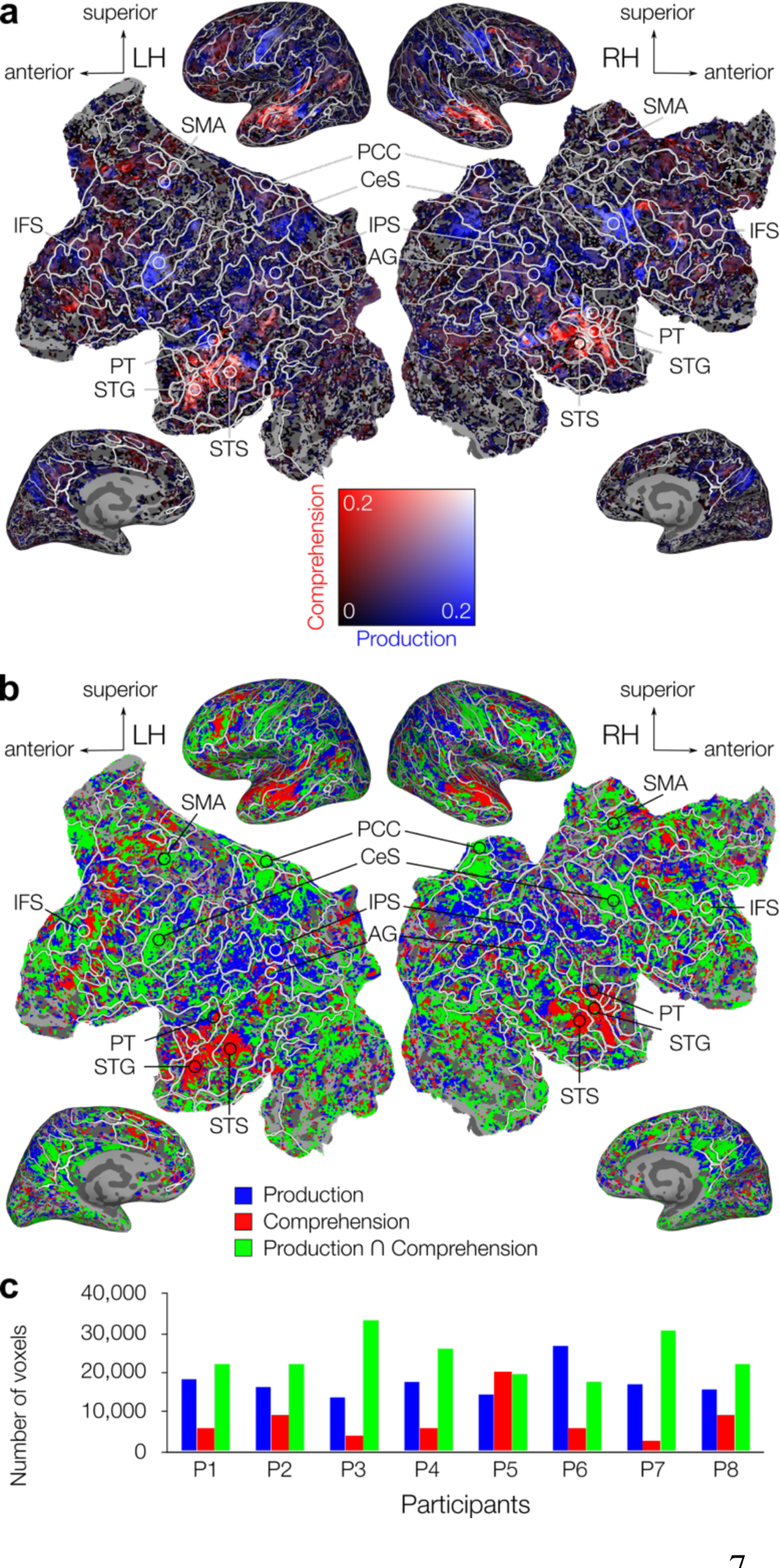
Linguistic selectivity for production and comprehension. **a**, Unique variance explained by production and comprehension in one participant (P8) are shown in blue and red, respectively. **Extended Data Fig. 4**summarizes results for all the participants. **b**, Spatial organization of bimodal and unimodal selective voxels. Green voxels best explained by the intersection of production and comprehension, while blue and red voxels are best explained by either production or comprehension. **Extended Data Fig. 5** summarizes results for all the participants. **c**, The number of voxels of the best variance partition.

## Linguistic weight correlation

We investigated the representational similarity between linguistic production and comprehension across the cortex. Previous research suggests that semantic and pragmatic information is integrated in amodal frontal, temporal, and parietal regions, including the semantic system^5,6,26,29^ and the default mode network^7,8^ (DMN), and that similar neural activity is involved in mutual understanding between narrative speaker and listener^1,2^. We hypothesized that similar linguistic representations in the amodal regions reflect mutual semantic and pragmatic information between language production and comprehension, allowing individuals to construct a coherent meaning. To examine representational similarity, we calculated the Pearson correlation coefficient between model weights for language production and comprehension (GPT-Neox, 2,816 dimensions) for each voxel within brain regions that showed significant predictions (**Fig. 1d**). We found positive correlations within the amodal regions (*P* < 0.05, two-sided, FDR corrected), with a cluster of high positive correlation in STG/STS (**Fig. 4a** and **Extended Data Fig. 6**). Conversely, a negative correlation was observed across the cortex (*P* < 0.05, two-sided, FDR corrected), with clusters of strong negative correlation in the motor cortex, including the ventral CeS and SMA (**Fig. 4b** and **Extended Data Fig. 7**). These results suggest that some voxels in the amodal regions encode meaning similarly across production and comprehension, supporting coherent linguistic information processing and enabling comprehension of language produced either by oneself or others.

**Fig. 4.**
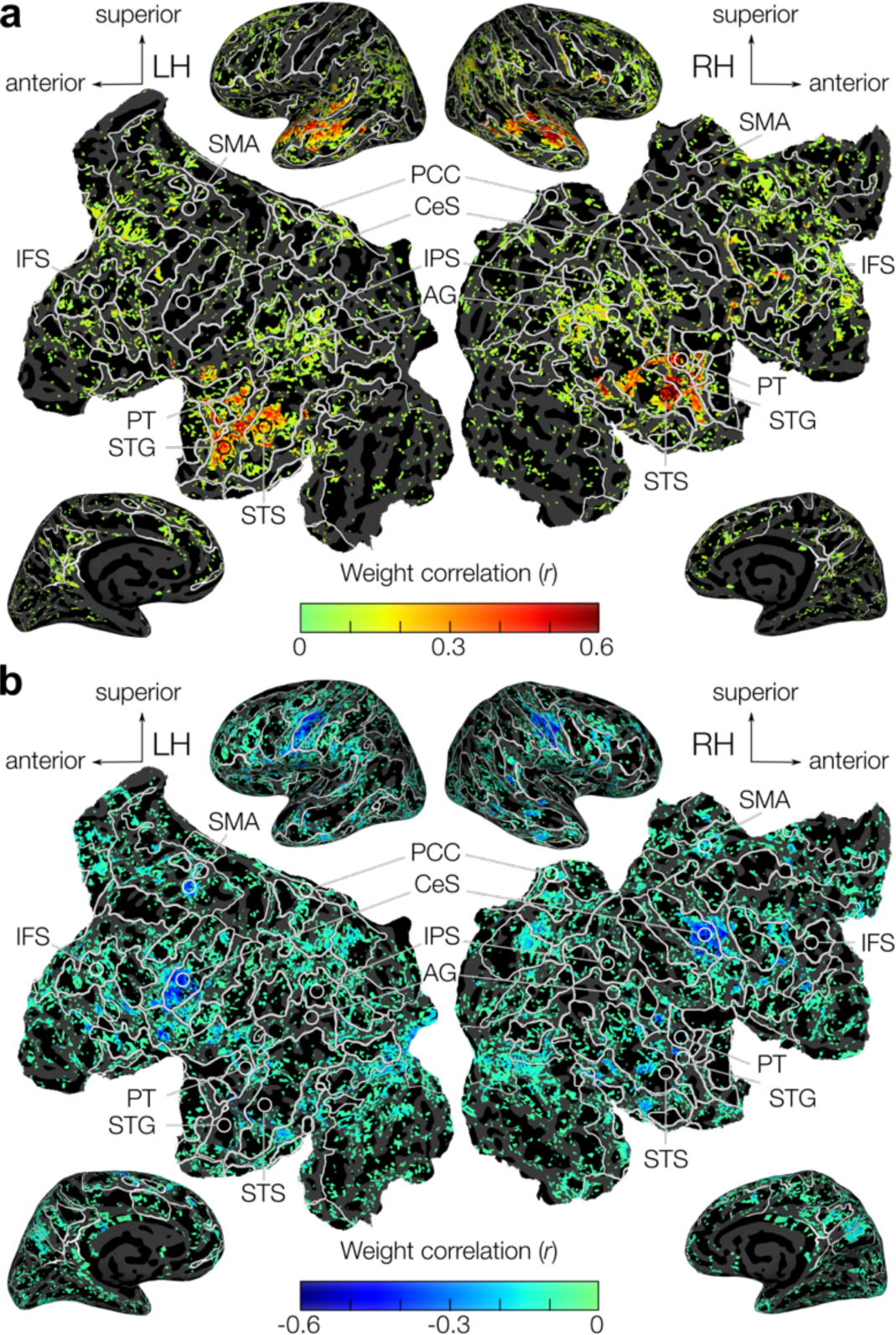
Weight correlation between linguistic production and comprehension. **a**, Positive correlations were observed in the amodal regions in frontal, temporal, and parietal cortices. **Extended Data Fig. 6** summarizes results for all the participants. **b**, Negative correlations were observed across the cortex. Correlations were calculated in voxels that showed significant predictions (**Fig. 1d**). **Extended Data Fig. 7** summarizes results for all the participants. Only voxels showing significant correlations are shown (*P*< 0.05, two-sided, FDR corrected).

## Principal components of representations

Finally, we sought to identify the specific content represented in each voxel. Given the high-dimensional data of tens of thousands of voxels, we used principal component analysis (PCA) to identify low-dimensional subspaces while retaining as much of the original variation as possible^5,37^. We applied PCA to the estimated linguistic weights aggregated across participants separately for production and comprehension, yielding 2,816 orthogonal dimensions for each modality. To determine whether each principal component (PC) explained significantly more variance than would be observed by chance, each PC was compared with the PC of the corresponding stimulus (i.e., utterance) matrices. Four and six PCs were significant for production and comprehension, respectively (**Fig. 5a, d**), suggesting that our fMRI data contain four and six dimensions for production and comprehension (**Extended Data Table 1**).

**Fig. 5.**
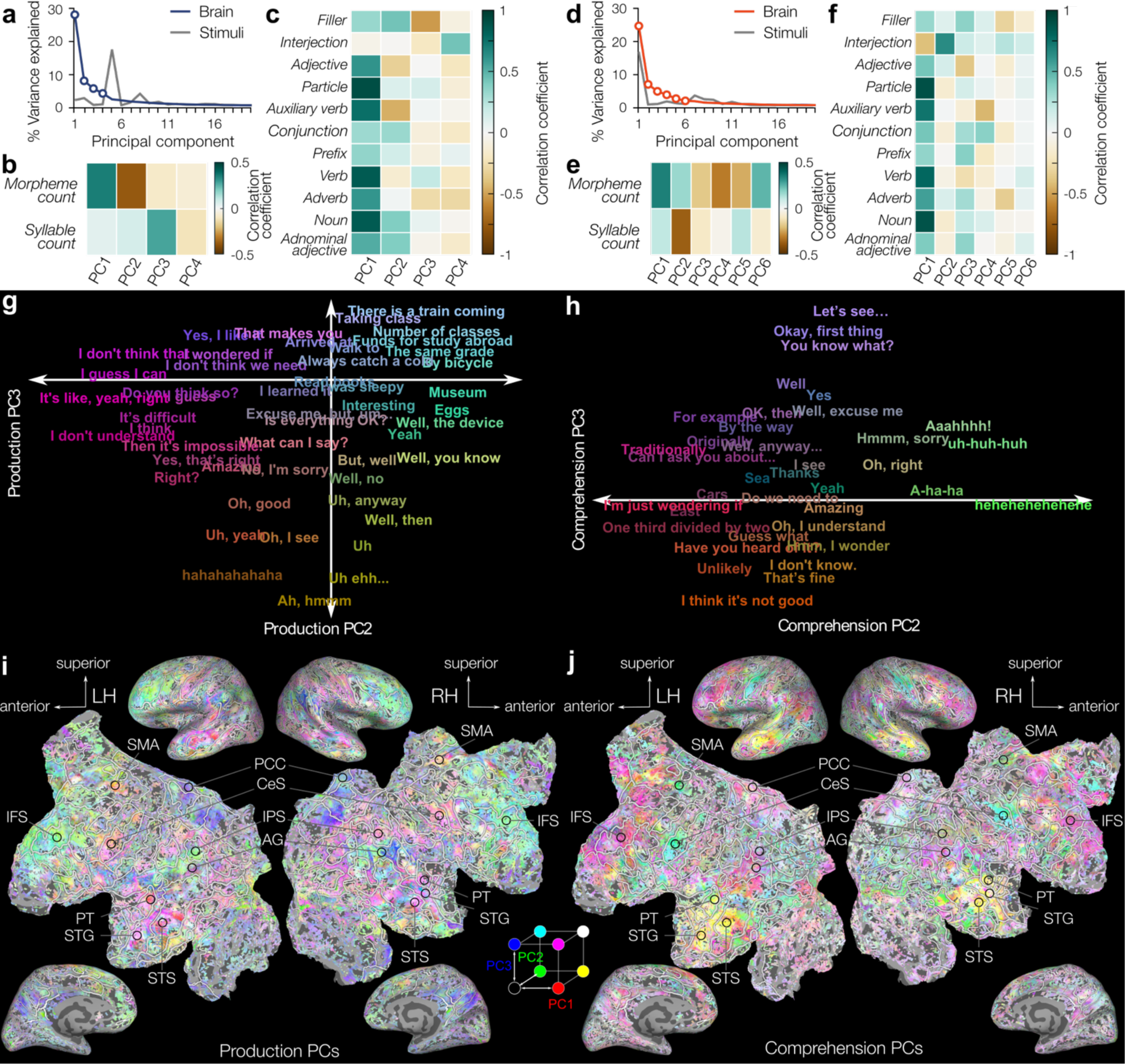
Principal components of voxel-wise linguistic models. Production and comprehension PCs are illustrated in left and right panels, respectively. **a/d**, Variance explained by the top 20 PCs for linguistic weights and utterance stimuli that are sorted by the stable matching algorithm. **b/e**, Partial Pearson correlation between low-level features and PC loading. **c/f**, Group mean Pearson correlation between regression weights for part-of-speech and PC scores averaged across the participants. **g/h**, Some of the utterances mapped in PC2 and PC3 are colored in RGB colormap. **i/j**, The significant three PCs (PC1, PC2, and PC3) are colored and mapped using RGB colormap. **Extended Data Fig. 8** and 9 summarizes results for all the participants.

To interpret the estimated low-dimensional space, we used various types of information. First, to visualize the production space and comprehension space, we projected the utterances in the dialogue experiment (total 25,854 and 25,272 utterances for production and comprehension, respectively) onto each dimension by calculating Pearson correlation between the linguistic embedding and each PC coefficient (**Supplementary Tables 1 –20**). Each utterance was assigned an RGB colour representing the red, blue and green channels of PC1, PC2, and PC3 (**Fig. 5 g, h**). Second, to investigate whether each dimension was involved in low-level linguistic properties, correlations were calculated as follows. To examine the complexity of an utterance in terms of syntax and phonology, partial Pearson correlations were calculated between the PC loading and the number of morphemes in the utterance controlling for the number of syllables and vice versa across the utterances (two-sided, FDR corrected, **Fig. 5 b, e**). Additionally, the Pearson correlation between each PC score and the individual part-of-speech weights estimated in the voxel-wise modelling was calculated to examine the word-level property within individual participants, and then the Pearson correlation coefficients were averaged across the participants (**Fig. 5 c, f**). Finally, we mapped the PC scores using the same RGB colour scheme to visualize the pattern of dialogue content selectivity across the cortex (**Fig. 5 i, j**, **Extended Data Fig. 8**, 9).

Then we examined each of the PCs in detail. It should be noted that some expressions were difficult to translate into English, and that the experiment and analysis were conducted in Japanese. First, both production and comprehension’s PC1 showed a negative correlation only with the weights of interjection across voxels concatenated across participants (**Fig. 5 c, f**), while also showing a strong positive correlation with the other part-of-speech weights. This suggests that the first dimensions represent turn-taking: speech production in the participant’s own turn and comprehension in the partner’s turn. Speech production in one’s own turn was involved in the speech motor control network (**Extended Data Fig. 10**), which includes CeS, PreCeG, SMA, PT, and MTG/STG. The primary and premotor cortex integrates acoustic-phonological and motor-articulatory processing^33^, and superior temporal regions are involved in auditory feedback control^31^. In contrast, comprehension in the partner’s turn involved several networks (**Extended Data Fig. 10**), including the auditory cortex, language network^38^ and the DMN, which includes the MPFC, PCC, TPJ, and STS. These regions are involved in the integration of extrinsic and intrinsic information and mentalizing when listening to a partner^8^.

Next, we examined PC2 to PC4 in terms of speech production. One end of PC2 favours proper nouns, especially places such as name of a university, with fewer morphemes (partial correlation *ρ* = −0.38, *P* < 0.0001, **Fig. 5b**). This is associated with the FPN, including the SFS/MFG, IFS/IFG, and IPS^30^, and with AG and MTG/ITG. These regions have been consistently linked with word retrieval^39^. The other end of PC2 is associated with utterances expressing opinions and mental states, such as ‘*I think*’, ‘*I would like to*’ (**Fig. 5g**), which relate to the DMN regions, including the PCC, TPJ, and STS. This finding is consistent with the DMN’s role in self-mentalizing^40^. In addition, one end of PC3 favours multi-word utterances, such as ‘*you basically say yes*’, ‘*be able to create a base*’, and ‘*Someone was next to me*’ (**Fig. 5g**), which contain many syllables (partial correlation *ρ* = 0.28, *P* < 0.0001). This is associated with the STS and TPJ, which are involved in syntactic sentence construction^41^. The other end of PC3 favours non-lexical fillers and laughter utterances such as ‘*Hmmm*…’, ‘*Well*…’, and ‘*Hahaha*’, that are idiosyncratically involved in brain regions across participants, over the association cortex in frontal, temporal, and parietal regions that have been implicated in increased activity during naturalistic filler production^11^. Finally, one end of PC4 favours backchannels such as ‘*yes*’ and ‘*right*’, associated with speech motor areas including CeS and SMA. The other end of PC4 favours utterances with demonstrative expressions, such as ‘*such*’ and ‘*something like that*’, which are involved in the dorsal visual pathway associated with processing of demonstratives^42^.

Next, we examined PC2 to PC6 in terms of speech comprehension. One end of PC2 is associated with a non-lexical backchannel and laughter utterances with fewer syllables (partial correlation *ρ* = −0.37, *P* < 0.0001, **Figure 5e**) that require little working memory load and involvement with the sensorimotor and auditory cortex that has been linked to involuntary laughter perception^43^. Conversely, the other end of the PC2 is related to utterances with many syllables that involve the FPN^30^ and require high working memory load. One end of the PC3 favours utterance-initial fillers^10^ with fewer morphemes (partial correlation *ρ* = −0.18, *P* < 0.0001), indicating that the conversational partner is attempting to take the floor, such as ‘*uh, well, then, (I have a little favour to ask you)*’ or starting a conversation such as ‘*okay then first*’ and ‘*you know what.*’ This is associated with the DMN, which is thought to form a context-dependent model of the unfolding situation^8^. Conversely, the other end of PC3 favours utterances that occur at the end of sentences such as interrogatives in Japanese (**Figure 5j**), and involves the language network^38^ that is associated with high constituent size in sentence construction^41^. Moreover, one end of the PC4 favours adverbial phrases with fewer morphemes (partial correlation *ρ* = −0.31, *P* < 0.0001), such as ‘*traditionally*’, ‘*originally*’, and ‘*now then*’, and involves regions that are less consistent across participants. The other end of PC4 favours utterances that request the partner to start speech production, such as ‘*I just wanted to ask you what you think*’, ‘*Do you understand?*’, and ‘*Have you heard?*’, and is involved with the DMN that is associated with thinking about oneself and others^40^ and with the speech planning network, including MFG, IFG, and PreCeS^9^. Furthermore, one end of the PC5 favours concrete nouns with fewer morphemes (partial correlation *ρ* = −0.23, *P* < 0.0001) such as ‘*a litre of milk*’, ‘*train*’, and ‘*mathematician*’, and is involved in the language network including the IPS, STS/MTG and IFS/IFG/PreCeS^38^. The other end of PC5 prefers non-lexical repetitive sounds of backchannels and laughter, such as ‘*hahahahaha*’ and ‘*ah ah ah.*’ This is involved in the DMN’s engagement in other-mentalizing about what the sound means – particularly for voluntary laughter^43^. Finally, one end of PC6 favours utterances with more morphemes (partial correlation *ρ* = 0.25, *P* < 0.0001), including a Japanese expression (‘*tteiu*’) that describes something the listener may not know. This is related to sensorimotor regions around the CeS, thought to generate speech sequence representations that are transformed into short-term memory^32^. The other end of PC6 favours utterances involving number and calculation, such as ‘*one third divided by two*’, ‘s*ix hundred thousand and one yen*’ and ‘*there are two thirds.*’ Consistent with previous studies on number/calculation processing^44,45^, PC6 is involved in IPS/AG and IFS/PreCeS. Taken together, these results suggest that successful dialogue is shaped by the coordination of linguistic and non-linguistic cognitive abilities and the complementary roles of production and comprehension, involving multiple networks, including the language network, speech-motor network, sensorimotor areas, auditory cortices, DMN, and FPN.

## Discussion

Our most striking finding was that the separate linguistic representations of production and comprehension are interwoven in the same networks. Notably, we observed similar linguistic representations in the amodal regions where conceptual semantic knowledge and social and pragmatic context are unified into meaning^5–8^. These separate and similar linguistic representations may contribute to making sense of language uttered by oneself or others, and the complementary adaptive interaction between self and others.

Our results provide a more detailed understanding of dialogue beyond the simple alignment account^13^, showing that two distinct speech representations reflect complementary roles between interlocutors. First, the correct control of vocal interaction was a more important factor in the speech representations than dialogue content, as evidenced by our finding that turn-taking (PC1 in both modalities) explained the largest amount of variance in the linguistic model weights. This is consistent with the empirical evidence that turn-taking is a universal system observed across languages^46^ and all the major primate clades^4^. Second, backchannels, which are non-content words that are used specifically in dialogue, are thought to facilitate smooth conversation rather than providing conceptual content^10^. We found that comprehension PC2 distinguished between low-syllable backchannels and high-syllable content words, suggesting that hearing backchannels does not interfere with speech production because it only requires minimal working memory load. We also found that production PC4 distinguished between backchannels and demonstrative expressions that point the listener’s attention to the previously spoken utterances, suggesting that backchannels signal understanding without disturbing the listener’s attention. Together, these findings provide unique neurobiological evidence that backchannels play a key role in smooth conversation by confirming shared understanding with minimal cognitive load. Finally, fillers are thought to serve several purposes, such as signalling a delay in the upcoming speech production and reducing the cognitive load for linguistic dual-tasking of speech planning while listening to the partner^10^. Consistent with this idea and a previous study^11^, we found that the production of fillers (production PC3) was associated with brain regions that were idiosyncratic across participants (e.g., MTG/ITG, IFS, SMA), suggesting that different cognitive processes are carried out, including lexical retrieval, selection, and speech suppression. We also found that utterance-initial fillers or words (comprehension PC3) were involved in the DMN, which integrates extrinsic upcoming information with intrinsic memory and knowledge^8^. These utterances may serve to introduce a new context and draw attention to the unfolding narrative. Thus, although fillers are devoid of conceptual content, they serve a pragmatic role that is specifically meaningful between the interlocutors depending on the context in which they are uttered. Taken together, our results highlight the important complementary roles of speech production and comprehension in real-life dialogue, as exemplified by turn-taking, backchannels, and fillers, and support the idea that successful dialogue is organized by interpersonal synergy^16,17^.

Our voxel-wise encoding model, combining a natural dialogue experiment with large language models, revealed the unique contribution of linguistic and non-linguistic abilities in everyday conversation. In agreement with recent studies on speech production/comprehension^34,35^, utterances from both speech production and comprehension (PC1 in both modalities) were consistently represented in the language network. Importantly, the language network represented linguistic abilities such as sentence construction^41^ (production PC3, comprehension PC3) and lexical retrieval/selection^39^ (production PC2, comprehension PC5), suggesting a specific role in linguistic processing.

In contrast, we identified distinct non-linguistic abilities that are essential for effectively conveying one’s own thoughts and inferring and understanding the thoughts of a partner. First, the DMN has been proposed as a sense-making network that integrates extrinsic information with intrinsic knowledge and memory^8^. Our results provide evidence that the DMN plays many important roles in dialogue, representing different social cognitive functions, including self-mentalizing^40^ when talking about oneself (production PC2) or being asked about oneself (comprehension PC4), other-mentalizing when listening to the partner’s turn^40^ (comprehension PC1) and when inferring the sounds and laughter that the partner vocalizes^43^ (comprehension PC5), and the context of the upcoming narrative that the partner unfolds^8^ (comprehension PC3).

Second, the FPN has been considered as an executive control network for cognitively demanding tasks^30^. Our results suggest that (part of) the FPN represents utterances in cognitively effortful situations^11^, including lexical retrieval/selection^39^ during the utterance of appropriate proper nouns (production PC2), often with the utterance of fillers (production PC3), and the processing of high-syllable utterances that burden working memory load^47^.

Third, the speech motor network, including the PreCeG/CeS, SMA and PT/MTG, is primarily involved in speech motor planning and execution through laryngeal motor control and auditory processing^33^. We found that this network is involved in speech production during one’s own turn (production PC1), and during the partner’s turn (backchannels, production PC4). Interestingly, this network was also involved in speech comprehension where it was shown to represent backchannels and involuntary laughter uttered by the partner (comprehension PC2), and adjectives asking the partner to answer the question (comprehension PC4). Our results provide evidence that the speech motor network is always involved in speech production and the comprehension of non-lexical sounds or utterances that invite the hearer to start talking.

Fourth, adjacent to this network, sensorimotor regions, including PostCeG/PostCeS, are thought to represent speech perception that is later transformed into working memory^32^. We found that the sensorimotor regions particularly represent utterances that seem unfamiliar to the participants (comprehension PC6).

Furthermore, the dorsal visual pathway has been implicated in the processing of demonstrative expressions^42^, although it is usually associated with visuospatial rather than language processing. We found that the dorsal pathway represents utterances that refer to previous discourse, such as ‘*something like that*’ in production PC4. Our results suggest that the dorsal pathway provides a unique means of communication based on pragmatic inference to refer to a specific part of the previous discourse in just a few non-content words.

Finally, number and calculation were represented in IPS/AG and IFS/PreCeS (comprehension PC6), which are hub regions for number and calculation processing^44,45^. This suggests that number and calculation are uniquely represented in the cortex as quantitative information that is distinct from language. These results emphasize that smooth conversation is enabled by both linguistic and extra-linguistic information, which depend on the coordination of the full range of cortical networks, from low-level sensorimotor to high-level language networks and the DMN.

The separate representations we have identified may play an important role in several cognitive abilities related to social communication, such as simultaneous speech planning and speech perception^9,15^, prediction of the partner’s speech^48^, and mental simulation of dialogue where one can think about what a partner would say if they said something^49^. An interesting possibility is that these separate representations may be acquired through a developmental process between infants and caregivers, providing the basis for the internalization of others and the simultaneous accommodation of multiple perspectives^50^. Future work is needed to gain further insights into the separate representations for speech production and comprehension.

A limitation of our experimental paradigm was that we could not rule out the possibility that our linguistic representations for production and comprehension reflect representations for self and other. To resolve this potential confound, future research could, for example, use a three-person dialogue to examine linguistic representations for self, other A, and other B, to better understand the production-comprehension and self-other separation. Different linguistic representations for other A and other B would imply that linguistic representations vary depending on the identity of the agent.

## Methods

### Participants

Eight healthy native Japanese speakers (P1–P8) participated in the fMRI experiment. P1 (male, age 22), P2 (male, age 22), P3 (male, age 23), P4 (female, age 22), P5 (male, age 20), P6 (female, age 20), P7 (female, age 20), and P8 (male, age 20). They were all right-handed according to the Edinburgh Handedness Inventory^51^ (with a laterality quotient score of 75–100) and had normal hearing and normal or corrected-to-normal vision. The Ethics and Safety Committee of the National Institute of Information and Communications Technology, Osaka, Japan, approved the experimental protocol, and all participants provided written informed consent.

### Natural dialogue experiment

The dialogue task involved chatting about 27 specific topics, such as self-introduction and favourite classes. These topics were selected to cover the semantic space in our daily lives and to reproduce many cognitive domains necessary for daily dialogue, including knowledge, recall, imagination, temporal and spatial cognition. As a reference resource, we used a corpus of everyday Japanese called the Corpus of Everyday Japanese (CEJC) available at https://www2.ninjal.ac.jp/conversation/cejc-monitor.html. Each fMRI run started with a specific topic and lasted 430 seconds. Participants were instructed to behave as they would in a normal dialogue in terms of content, turns, and topic changes. They listened to the interlocutor’s voice through insert earphones designed for fMRI research, and their recorded speech was played back to the interlocutor through a noise-cancelling microphone. The speech of participants and interlocutor were recorded separately by computer. Each participant was scanned for 27 runs over the course of four sessions in all but three sessions for one participant (P3). As only one run was correctly collected within one session, the remaining data collected in three sessions were used in the analysis for participants P2 and P5. In each session, two to ten runs of data was used for the analysis (**Extended Data Table 2**). The fMRI volumes where participants produced an utterance (mean ± SD = 5,758.1 ± 863.5, range 4,255–7,057; 50.4 ± 6.4%, 39.6–60.8%) and comprehended an utterance (mean ± SD = 5,667.9 ± 408.3, range 5,212–6,296; 49.7 ± 2.9%, 46.2–54.2%) were comparable (**Supplementary** Fig. 1).

### MRI data acquisition

MRI data were collected on a 3T MRI scanner (Siemens MAGNETOM Prisma for P1-P5, and Siemens MAGNETOM Prisma Fit for P6-P8) at CiNet using a 64-channel head coil. Functional scans were collected using a T2-weighted gradient echo multi-band echo-planar imaging (EPI) sequence^52^ in interleaved order covering the entire brain with repetition time (TR) = 1.0 s, echo time (TE) = 30 ms, flip angle = 60 deg, matrix size = 96 x 96, field of view = 192 mm x 192 mm, voxel size = 2 mm x 2 mm x 2 mm, slice gap = 0 mm, 72 axial slices, multi-band factor = 6. Anatomical data were collected using a T1-weighted MPRAGE sequence with TR = 2.53 s, TE = 3.26 ms, flip angle = 9 deg, matrix size = 256 x 256, field of view = 256 mm x 256 mm, voxel size = 1.0 mm x 1.0 mm x 1.0 mm.

### Transcription

Dialogues were first transcribed at the morphological level using Microsoft Azure Speech To Text. The transcripts were corrected manually. Then one or more morphemes were concatenated into semantic-aware chunks with a duration around the fMRI TR (i.e., 1,000 ms). Finally, each chunk was time-aligned to the corresponding fMRI volume using the middle of the duration.

### Linguistic model construction

To model linguistic content, we used an open source pre-trained conversational large language model, GPT-Neox (https://huggingface.co/rinna/japanese-gpt-neox-3.6b-instruction-sft). Thismodel is based on an open source variant of GPT^21^, GPT-Neox^53^, and has been further fine-tuned to serve as an instruction-following conversational agent. Specifically, this model was first pre-trained to predict the next word from the previous context on 312.5 billion tokens from Japanese text datasets, Japanese CC-100 (http://data.statmt.org/cc-100/ja.txt.xz), Japanese C4 (https://huggingface.co/datasets/mc4), and Japanese Wikipedia (https://dumps.wikimedia.org/other/cirrussearch). This model was then fine-tuned to serve as a chatbot. The fine-tuning was performed using a subset of datasets translated into Japanese, Anthropic HH RLHF data (https://huggingface.co/datasets/Anthropic/hh-rlhf), FLAN Instruction Tuning data (https://github.com/google-research/FLAN), and Stanford Human Preferences Dataset (https://huggingface.co/datasets/stanfordnlp/SHP). The pre-trained model consists of 36 layers and 2,816 dimensions of hidden units. We focused only on layer 27 because previous studies reported that features from middle layers predict brain activity better than other layers^27,54^.

### Spectrogram model construction

Based on previous studies of language encoding models^25,27^, we used the Lyon’s passive cochlea model (https://github.com/theunissenlab/tlab/tree/master/AuditoryToolbox) to model the spectral feature space. This cochleagram uses approximately logarithmically spaced filters with a bandwidth given by

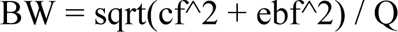

where cf is the characteristic frequency, ebf is the ear break frequency parameter set at 1,000 Hz, and Q is the quality factor set at eight. The output of this cochlear filter bank consisted of 80 waveforms between 264 and 7,630 Hz, spaced at 25% of the bandwidth. For the Nyquist frequency of the fMRI (0.5 Hz), a low-pass filter and then a Lanczos filter were applied.

### Part-of-speech, morpheme count, and syllable count model construction

We performed a morphological analysis of each utterance using MeCab (https://taku910.github.io/mecab/). MeCab estimates a part-of-speech for each morpheme from 11 categories: filler, interjection, adjective, particle, auxiliary verb, conjunction, prefix, verb, adverb, noun, and adnominal. We counted the number of morphemes and syllables in each utterance.

### Head motion model construction

To account for response variance correlated with head motion, we used the six motion parameters estimated in the motion correction pre-processing described below.

### fMRI data preprocessing

Motion corrections were performed for each run using the Statistical Parametric Mapping Toolbox (SPM8). All volumes were aligned using the first echo-planar imaging result for each participant. Low-frequency drift was removed using a median filter with a 120-s window. The response for each voxel was then normalized by subtracting the mean response and scaling the response to the unit variance. We used FreeSurfer^55,56^ to identify cortical surfaces based on anatomical data and register them with voxels for functional data. For each participant, the voxels identified in the cerebral cortex were used in the analysis (64,072–72,018 voxels per participant).

### Voxel-wise model estimation and testing

We used a finite impulse response (FIR) model to fit cortical activity measured in each voxel, capturing slow hemodynamic responses and their coupling to neural activity^23,57^. Although many fMRI studies use the canonical hemodynamic response function, this function assumes that the hemodynamic response function has the same shape across all cortical voxels. This can lead to an inaccurate modelling of brain activity because there is variation in hemodynamic responses across cortical regions^58^. Our FIR model consisted of five feature spaces (linguistic, part-of-speech, morpheme count, syllable count, and spectrogram) for production and comprehension and head motion. To capture hemodynamic delay, we concatenated the 5,822 features delayed by two to seven samples (2, 3, …, 7 s), yielding a total of 34,932 features. BOLD responses were modelled by multiplying the features by regression weights. We used an L2-regularized linear regression to estimate the 34,932 weights using training data. The optimal regularization parameter was evaluated using 10-fold cross-validation, with the 18 different regularization parameters ranging from 10 to 10 x 2^17^. We performed leave-one-session-out cross-validation, where one session of data was left out for testing and the remaining data was used for training. Prediction accuracy was calculated using Pearson’s correlation coefficient between the predicted signal and the measured signal in the test data. Statistical significance (one-sided) was tested by comparing the null distribution of correlations between two Gaussian random vectors with the same length as the test data. Estimated *P*-values were corrected for multiple comparisons using the false discovery rate (FDR) procedure^59^.

### Variance partitioning

To examine the unique variance explained by each set of features, we conducted variance partitioning, following similar approaches to prior studies that used voxel-wise modelling with multiple features^25,27^. First, the full sets of features were concatenated, and variance explained by the model was calculated (R^2^full). Second, to calculate unique variance explained by a set of features (e.g., part-of-speech in speech production), variance explained by a reduced model that excluded these features of interest from the full model was calculated (R^2^reduced). The unique variance explained by these features of interest was calculated using the following formula:

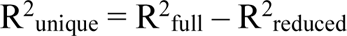

To ensure compatibility with the correlation coefficient, we used the square root (Runique) to represent the relative complement (as shown in **Figure 3a**).

### Principal component analysis

To identify a low-dimensional space from the estimated linguistic model weights for production and comprehension, we performed principal component analysis (PCA). We included only the 10,000 best predictive voxels from each participant to exclude noise from less predictive voxels. To remove temporal information, we averaged across the six delays for each feature. We then applied PCA to the model weights (2,816 dimensions) for production and comprehension separately, after concatenating voxels across all participants (2,816 dimensions x 80,000 voxels). To determine which weighted PCs explained significantly more variance than would be expected by chance, we compared the weighted PCs with the stimulus PCs. Specifically, we calculated confidence intervals of variance explained in each PC by bootstrapping the weighted PCA and stimulus PCA 1,000 times. To increase correspondence between the weighted PC and the stimulus PC, we used the Gale-Shapley stable marriage algorithm^5,37^.

### Region of interest abbreviations

Central sulcus (CeS), precentral sulcus/gyrus (PreCeG/PreCeS), postcentral sulcus/gyrus (PostCeG/PostCeS), supplementary motor area (SMA), inferior frontal sulcus (IFS), triangular part of the inferior frontal gyrus (IFG-Tri), opercular part of the inferior frontal gyrus (IFG-Op), middle frontal gyrus/sulcus (MFG/MFS), superior frontal gyrus/sulcus (SFG/SFS), inferior temporal gyrus/sulcus (ITG/ITS), middle temporal gyrus/sulcus (MTG/MTS), superior temporal gyrus/sulcus (STG/STS), planum temporale (PT), supramarginal gyrus (SMG), angular gyrus (AG), temporoparietal junction (TPJ), intraparietal sulcus (IPS), medial prefrontal cortex (MPFC), posterior cingulate cortex (PCC), superior parietal gyrus (SPG), anterior cingulate cortex (ACC), subparietal sulcus (SubPS), precuneus (PreC), parieto-occipical sulcus (POS), middle occipital sulcus (MOS), superior occipital sulcus (SOS).

## Data availability

All data used in the current study will be made publicly available at OpenNeuro before publication.

## Code availability

The MATLAB code used in the current study will be made publicly available at GitHub before publication.

## Acknowledgements

We thank JST ERATO JPMJER1801, CREST JPMJCR18A5, and MIRAI JPMJMI19D1 (for S.N.) for the financial support of this study.

## Author contributions

S.N. conceptualized the experiment. K.R. and S.N. designed the experiment. K.R. performed the experiment. M.Y., K.R. and S.N. analyzed and interpreted the results. M.Y. and S.N. wrote the paper. S.N. supervised the study.

## Competing interests

The authors declare no competing financial interests.

## Supplementary information

Supplementary Fig. 1 and Tables 1–20.

**Correspondence and requests for materials** should be addressed to Shinji Nishimoto.

## Extended Data

**Extended Data Fig. 1:**
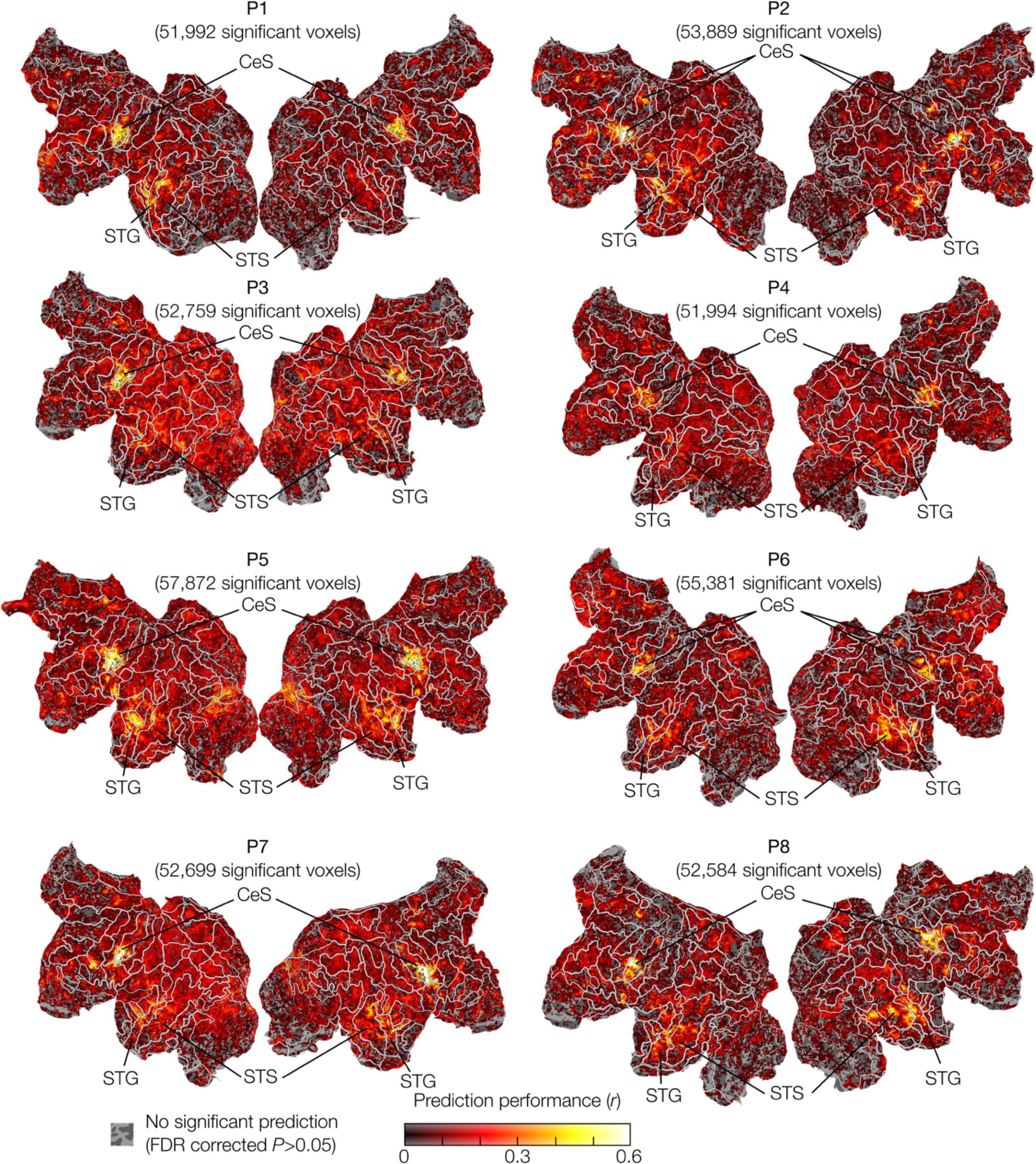
Voxel-wise model prediction performance. Prediction was performed using the linguistic (GPT-Neox) production and comprehension features and other features were discarded.

**Extended Data Fig. 2:**
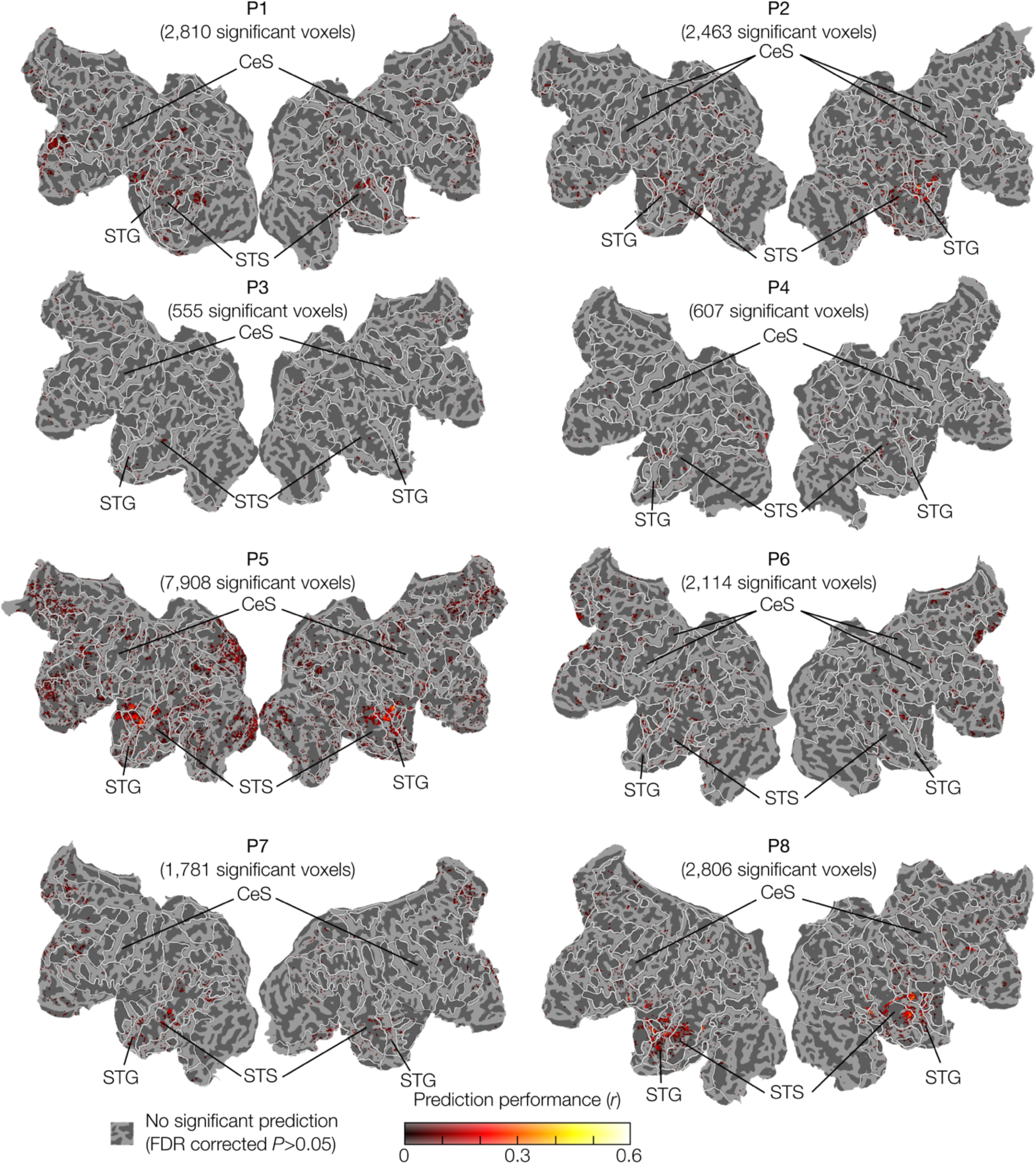
Voxel-wise model cross-modal prediction performance. Prediction was performed using the linguistic (GPT-Neox) production and comprehension features and other features were discarded.

**Extended Data Fig. 3:**
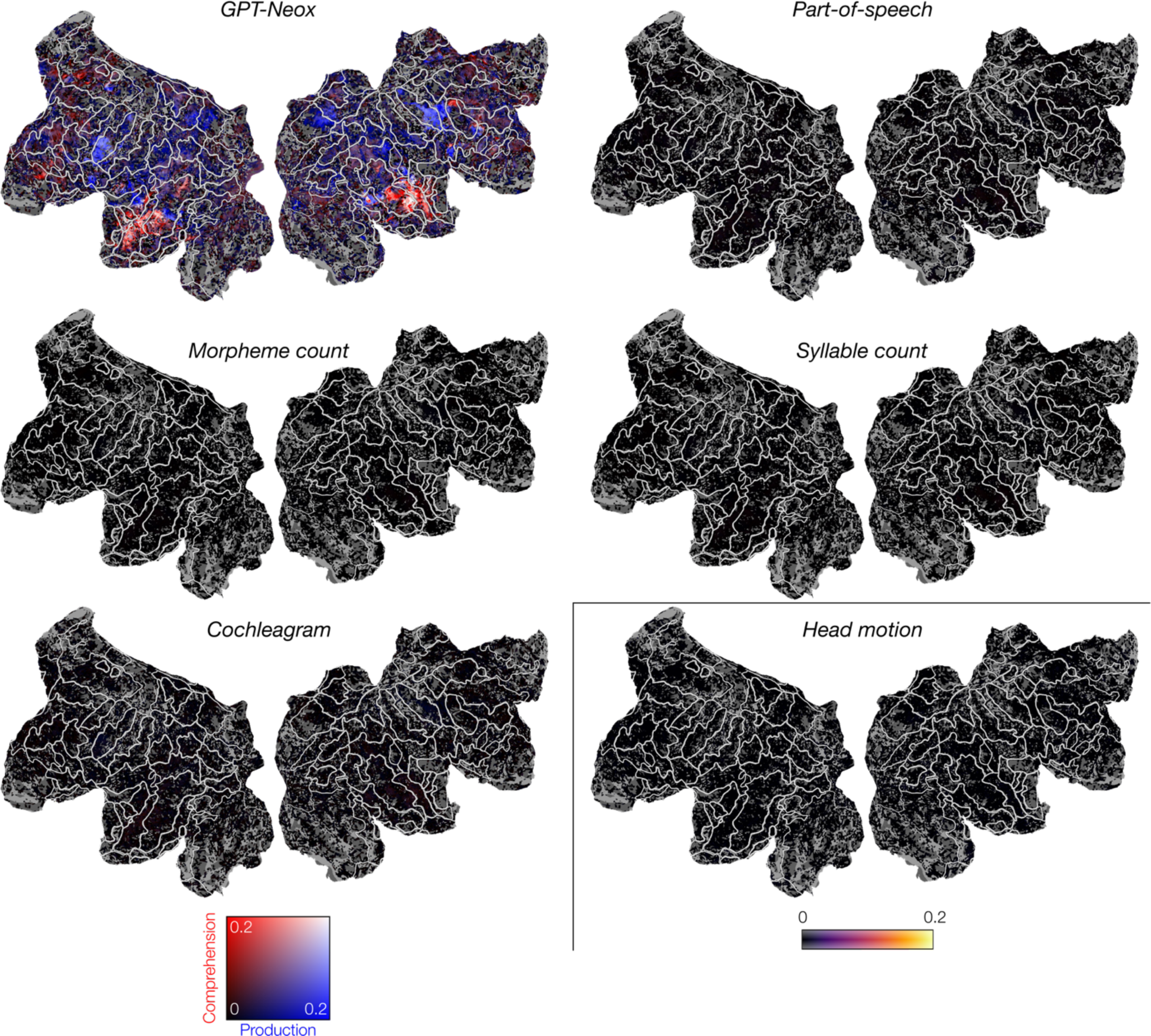
Unique variance explained by each set of features using the full model in one participant (P8).

**Extended Data Fig. 4:**
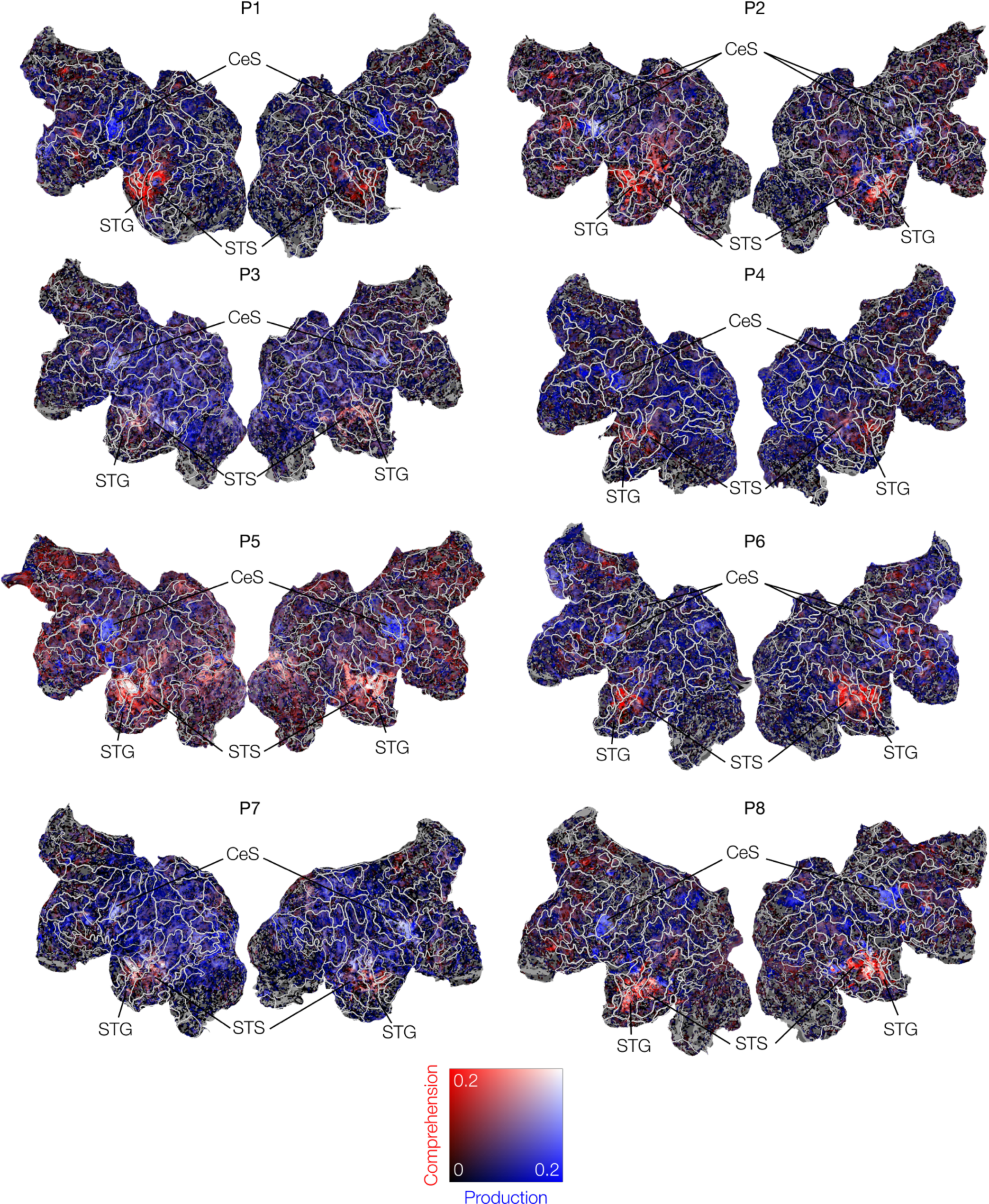
Unique variance explained by linguistic production or comprehension using the reduced model.

**Extended Data Fig. 5:**
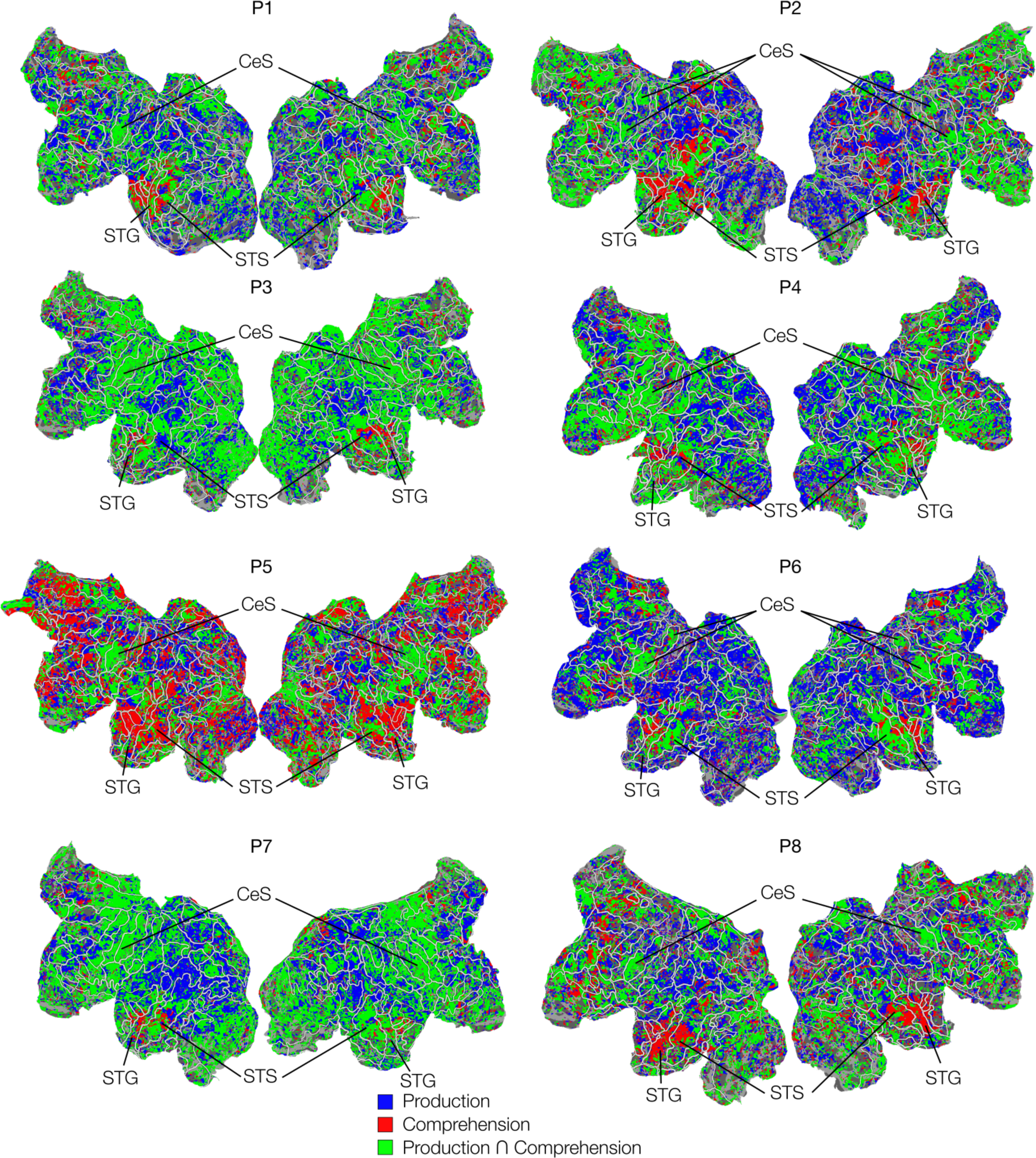
The best variance partition.

**Extended Data Fig. 6:**
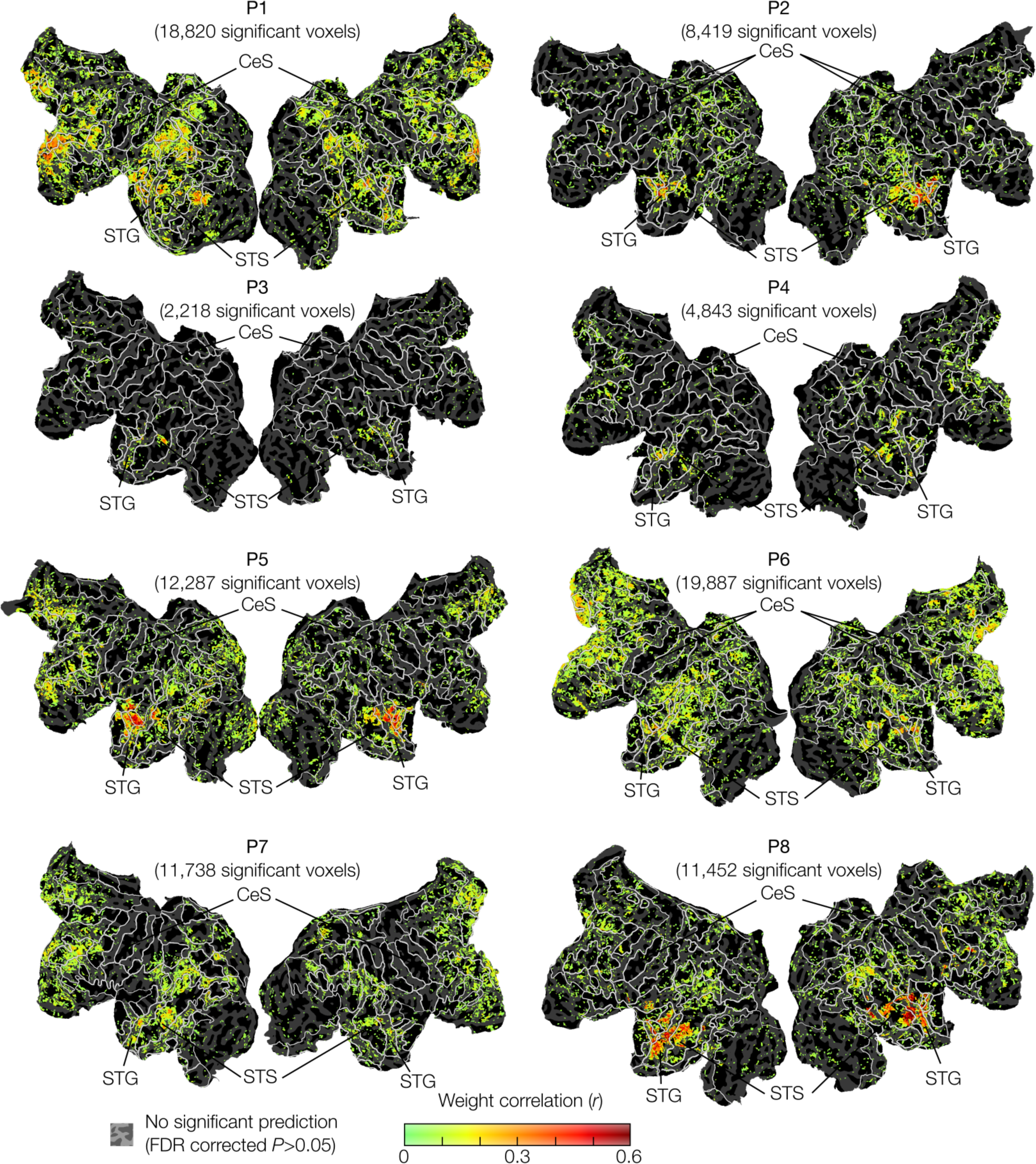
Significantly positive weight correlation between linguistic production and comprehension. The correlations were calculated within the voxels that showed significant same-modal prediction.

**Extended Data Fig. 7:**
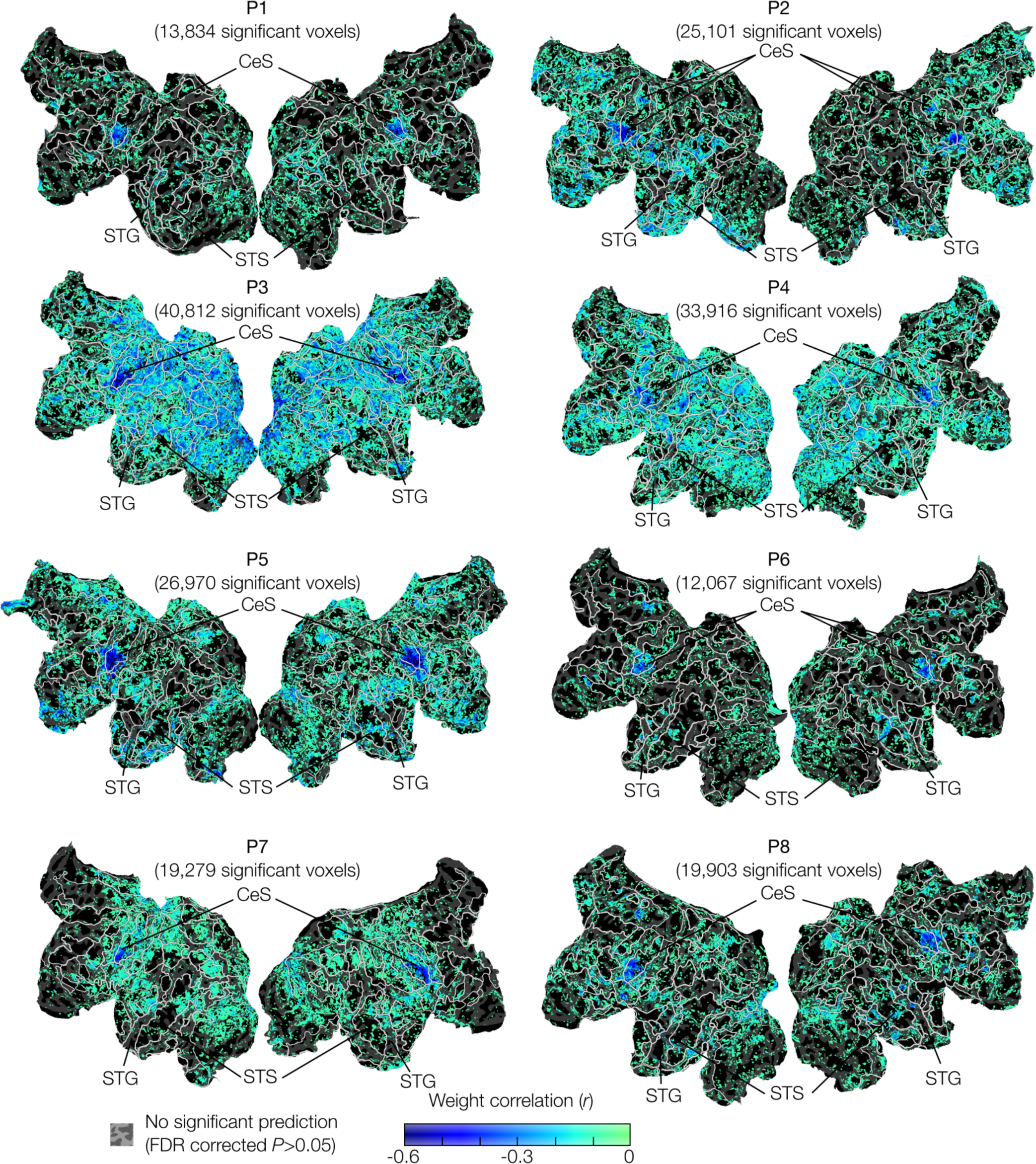
Significantly negative weight correlation between linguistic production and comprehension. The correlations were calculated within the voxels that showed significant same-modal prediction.

**Extended Data Fig. 8:**
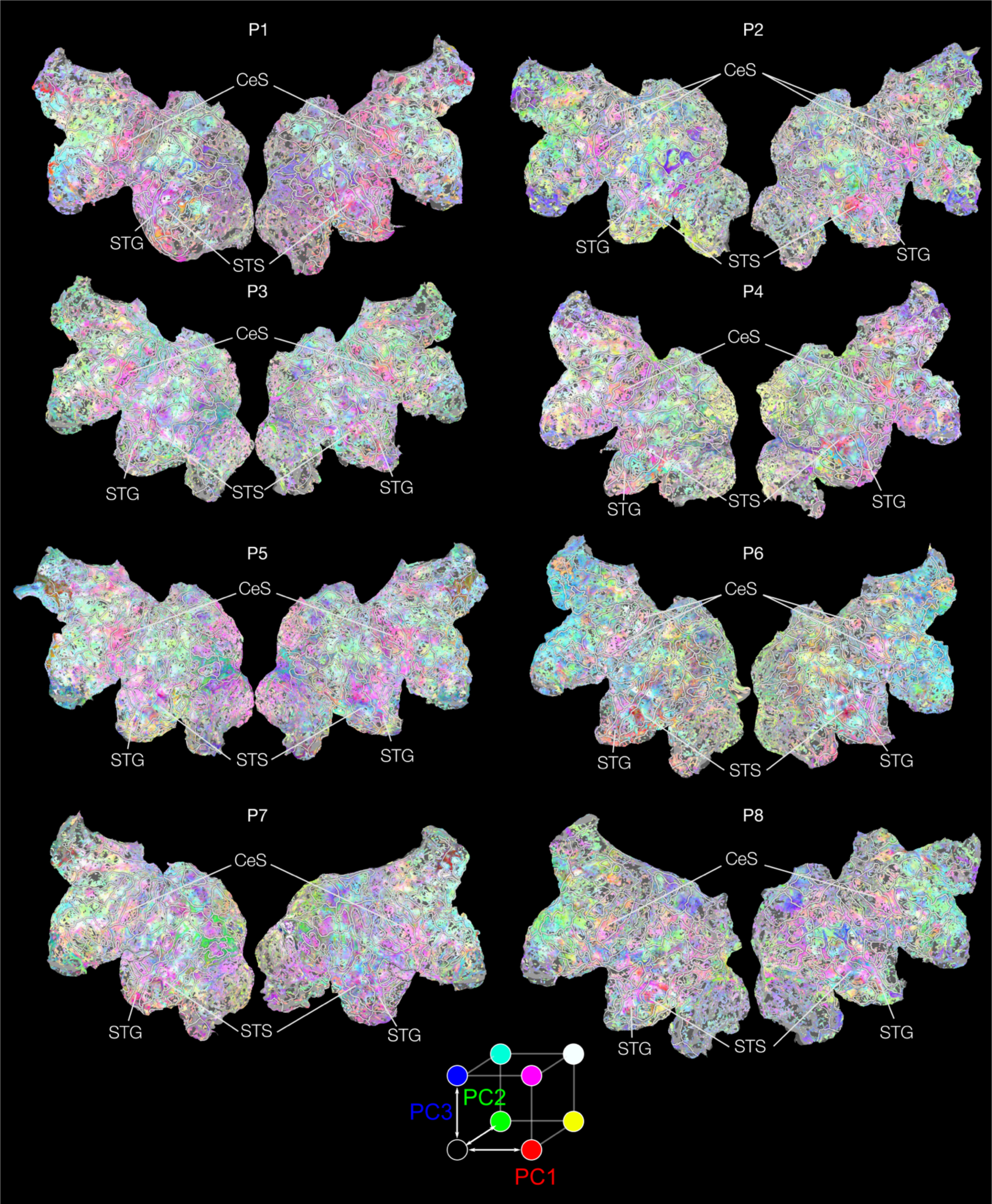
Principal components of linguistic production models. The principal component analysis (PCA) was performed within the top 10,000 predictive voxels, and then mapped onto the voxels that showed significant same-modal prediction.

**Extended Data Fig. 9:**
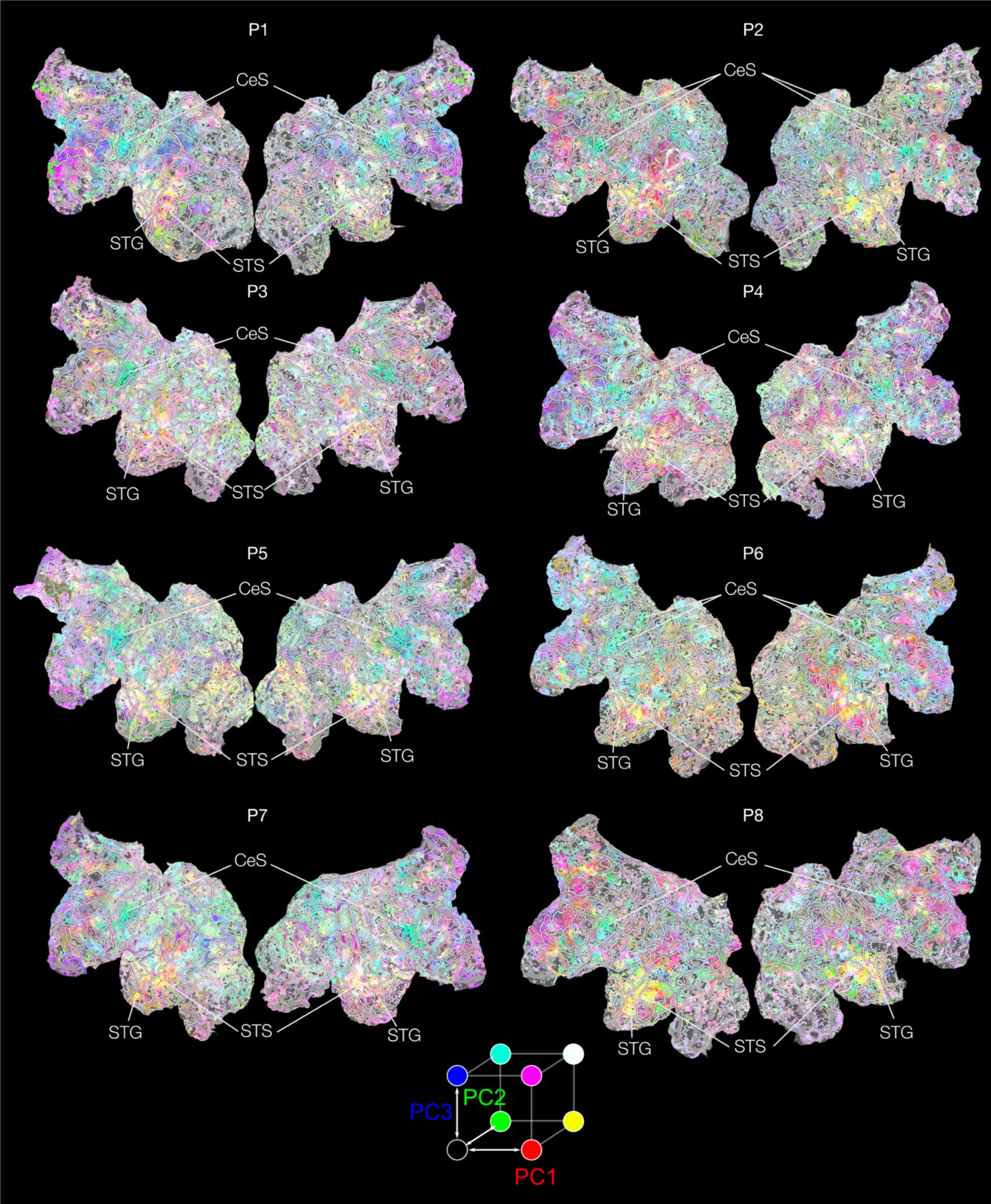
Principal components of linguistic comprehension models. The principal component analysis (PCA) was performed within the top 10,000 predictive voxels, and then mapped onto the voxels that showed significant same-modal prediction.

**Extended Data Fig. 10:**
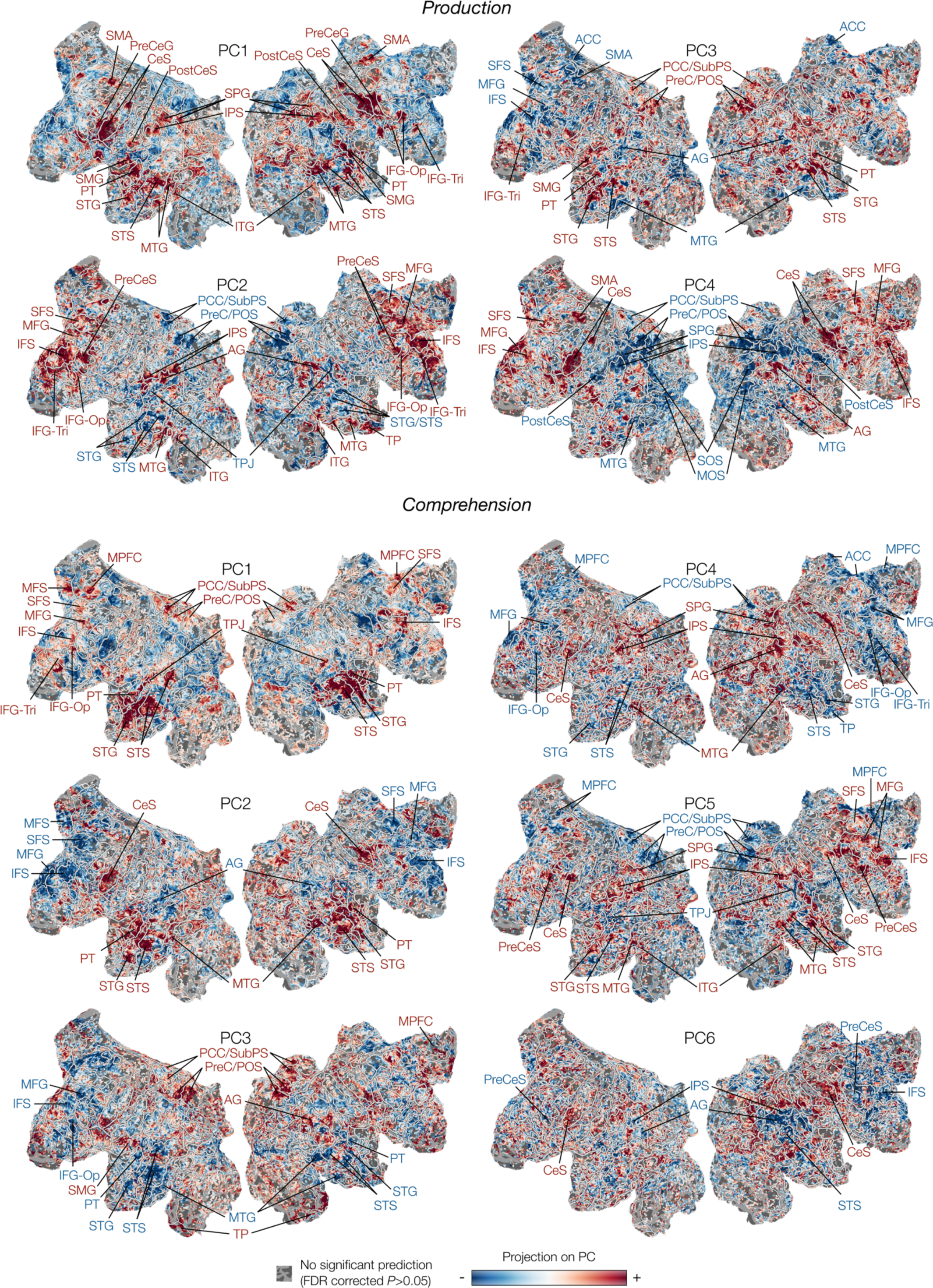
Principal components of linguistic models in one participant (P8).

**Extended Data Table 1.**
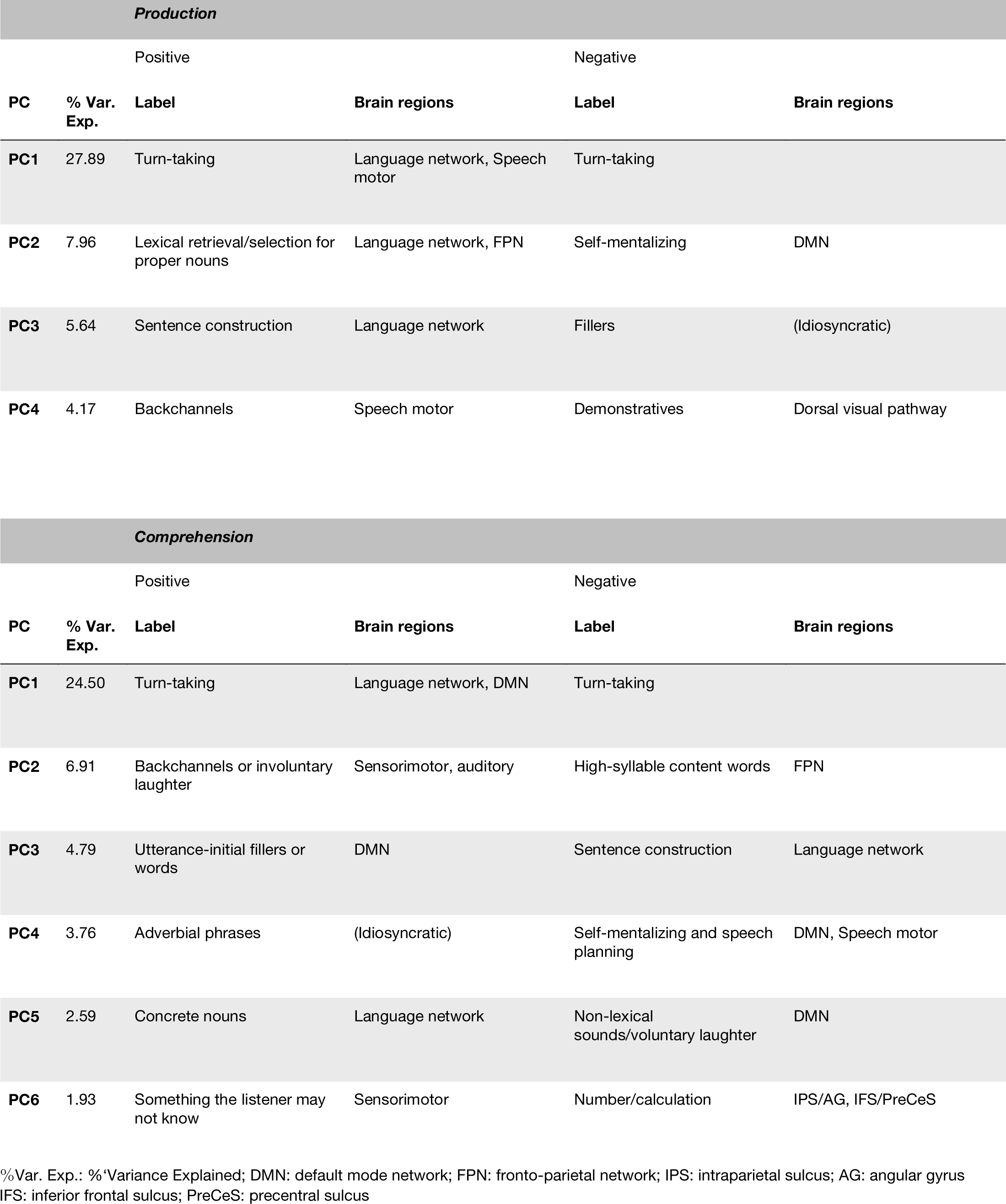
% Variance explained, labels, and relevant brain regions for each principal component.

**Extended Data Table 2.** The number of fMRI data samples [runs] in each session.

| **Participant** | **Ses. 1** | **Ses. 2** | **Ses. 3** | **Ses. 4** |
| --- | --- | --- | --- | --- |
| **P1** | 2,580 [6] | 3,870 [9] | 1,290 [3] | 3,870 [9] |
| **P2** | 3,010 [7] | 3,870 [9] | 3,870 [9] | N/A |
| **P3** | 3,440 [8] | 3,870 [9] | 4,300 [10] | N/A |
| **P4** | 3,010 [7] | 3,440 [8] | 3,870 [9] | 1,290 [3] |
| **P5** | 3,010 [7] | 3,870 [9] | 3,870 [9] | N/A |
| **P6** | 3,010 [7] | 3,440 [8] | 3,870 [9] | 1,290 [3] |
| **P7** | 3,010 [7] | 3,870 [9] | 3,010 [7] | 1,720 [4] |
| **P8** | 3,010 [7] | 3,870 [9] | 3,870 [9] | 860 [2] |

## Notes

### Competing Interest Statement

The authors have declared no competing interest.

