## Supplementary Information for "Cortical representations of languages during natural dialogue"

**Table of contents:**

Figure 1. The number of fMRI volumes that included utterances of participants (top) and the experimenter (bottom).

Table 1. Production PC1 highest correlation utterances (Turn-taking).
Table 2. Production PC1 lowest correlation utterances (Turn-taking).
Table 3. Production PC2 highest correlation utterances (Lexical retrieval/selection for proper nouns).
Table 4. Production PC2 lowest correlation utterances (Self-mentalizing).
Table 5. Production PC3 highest correlation utterances (Sentence construction).
Table 6. Production PC3 lowest correlation utterances (Fillers).
Table 7. Production PC4 highest correlation utterances (Backchannels).
Table 8. Production PC4 lowest correlation utterances (Ambiguous expressions).
Table 9. Comprehension PC1 highest correlation utterances (Turn-taking).
Table 10. Comprehension PC1 lowest correlation utterances (Turn-taking).
Table 11. Comprehension PC2 highest correlation utterances (Backchannels or involuntary laughter).
Table 12. Comprehension PC2 lowest correlation utterances (High-syllable content words).
Table 13. Comprehension PC3 highest correlation utterances (Utterance-initial fillers or words).
Table 14. Comprehension PC3 lowest correlation utterances (Sentence construction).
Table 15. Comprehension PC4 highest correlation utterances (Adverbial phrases).
Table 16. Comprehension PC4 lowest correlation utterances (Self-mentalizing).
Table 17. Comprehension PC5 highest correlation utterances (Concrete nouns).
Table 18. Comprehension PC5 lowest correlation utterances (Non-lexical sounds/voluntary laughter).
Table 19. Comprehension PC6 highest correlation utterances (Something the listener may not know).
Table 20. Comprehension PC6 lowest correlation utterances (Number/calculation).

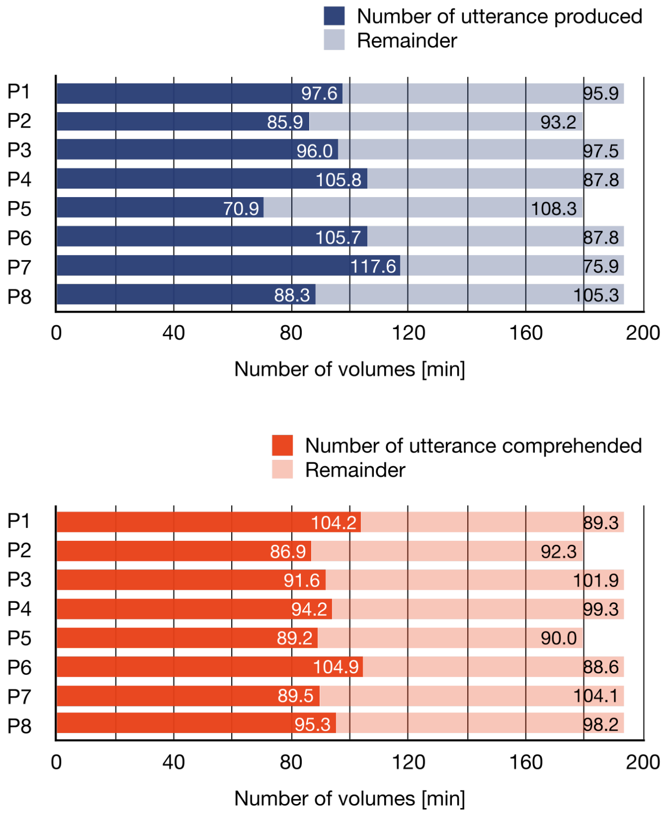


**Supplementary Fig. 1. The number of fMRI volumes that included utterances of participants (top) and the experimenter (bottom).**

### Supplementary Tables

To illustrate how utterances were related to the principal components (PC), each utterance was projected onto each PC by correlating the PC coefficients and the GPT embeddings. The top 20 most correlated utterances are included in the tables. The Japanese-English translation was performed by DeepL (https://www.deepl.com/translator), including the preceding and following context, and only the relevant utterances are shown in ***bold italic***. Non-lexical words, such as backchannels, fillers, and laughter, were manually modified by considering their sounds.

**Supplementary Table 1. Production PC1 highest correlation utterances (Turn-taking).**

| Rank | Correlation coefficient | Utterance |
|---|---|---|
| 1 | 0.165896 | My friends and I were ***talking about*** the meaning of this assignment. |
| 2 | 0.164770 | Someone was trying to figure out what the motivation ***would be*** to go into the store. |
| 3 | 0.164689 | I thought ***I should talk*** a lot. |
| 4 | 0.164581 | If everyone had the same face, ***it probably wouldn't*** feel so strange. |
| 5 | 0.163899 | how many cranes or turtles ***there are*** |
| 6 | 0.163811 | The main materials used are ***things like*** carbon and so on. |
| 7 | 0.163254 | ***It's just that*** my attitude ***changes***. |
| 8 | 0.163171 | ***I kind of hate*** that one. |
| 9 | 0.163153 | ***What do you mean?*** |
| 10 | 0.162327 | I'd rather have hands-on experience and that kind of thing ***than learn*** from books. |
| 11 | 0.162261 | ***I think*** looking at people's faces ***is losing*** its appeal. |
| 12 | 0.161871 | I was trying to see if it would be stimulating or not, but I guess ***that's the kind of*** project it is. |
| 13 | 0.161836 | But if it were cranes and turtles, ***it would be like*** how many cranes and turtles ***are there*** together. |
| 14 | 0.161795 | I'm a little nervous about this experiment, like ***I'm not sure if*** it's going to be useful. |
| 15 | 0.161654 | The best form of relationship is ***to start*** expanding one-on-one or something like that. |
| 16 | 0.161441 | I was trying to wake myself up by twisting my body ***and so on***. |
| 17 | 0.161412 | I was pretty sure that if I wasn't allowed to move, ***I would probably*** fall asleep. |
| 18 | 0.160949 | You should just make sure ***that*** appearance has something to do with sales before you make a decision. |
| 19 | 0.160869 | ***I don't know what*** a comfortable line is for me. |
| 20 | 0.160779 | I was just wondering ***what*** the research ***was like.*** |

**Supplementary Table 2. Production PC1 lowest correlation utterances (Turn-taking).**

| Rank | Correlation coefficient | Utterance |
|---|---|---|
| **1** | 0.105681 | ***Get, get*** married, ummm, would you like to... |
| **2** | 0.108919 | ***At, at*** [Train station] |
| **3** | 0.109514 | ***Consumer*** |
| **4** | 0.109594 | It's better to think about ***the question first.*** |
| **5** | 0.110112 | so, ***well, yeah***, yeah. |
| **6** | 0.110484 | Well, in the vicinity, I fish, ***well, in [Place name].*** |
| **7** | 0.110648 | He trained and trained and ***trained, and*** he was tired of doing it, so he |
| **8** | 0.110716 | ***And the other*** one is quite a swarthy-looking guy. |
| **9** | 0.111200 | [Railway] is, well, about ***north-north***east. |
| **10** | 0.111284 | [Railway] is, well, about north-north***east***. |
| **11** | 0.111426 | I believe the Department of Nursing is ***west*** of [Restaurant name] |
| **12** | 0.111467 | ***Two***, Russian taken as a second foreign language. |
| **13** | 0.111642 | If the other ***members*** of the team don't move. |
| **14** | 0.111690 | Dogs and rabbits ***and,*** as a separate category, |
| **15** | 0.111822 | Well, ***the point*** is, the point is, um... |
| **16** | 0.111920 | ***Father, mother, brother***, and me |
| **17** | 0.111929 | And divide into milk, ***bread and eggs.*** |
| **18** | 0.111931 | It's like an egg on top of that, ***and bread on top.*** |
| **19** | 0.111939 | There is a ***[Railway]*** line**.** |
| **20** | 0.111979 | ***Eh, heh.*** |

**Supplementary Table 3. Production PC2 highest correlation utterances (Lexical retrieval/selection for proper nouns).**

| Rank | Correlation coefficient | Utterance |
|---|---|---|
| **1** | 0.047039 | ***Fourth*** year student, ***Faculty of [Faculty name]***, [University name] |
| **2** | 0.042527 | I belong to the Comprehensive Psychology Department ***at [University name]***. |
| **3** | 0.042332 | It's called the Department of [Department] ***in the Faculty of [Faculty name]***. |
| **4** | 0.040928 | Some people ***in the ice hockey team.*** |
| **5** | 0.040910 | ***[University name]*** |
| **6** | 0.040299 | ***[University name]*** |
| **7** | 0.039816 | ***you know, in universities,*** in laboratories, where they put the birds. |
| **8** | 0.039803 | ***Engineering*** professors think it's a good idea. |
| **9** | 0.039703 | ***Faculty of [Faculty name]*** and [Library] Library straight down the road is |
| **10** | 0.039211 | ***[University name]*** |
| **11** | 0.039095 | ***[Place name]*** in [Place name] ***and*** also in [Place name], [Place name] Prefecture. |
| **12** | 0.039022 | When I have time, I go to ***the Faculty of [Faculty name] cafeteria***. |
| **13** | 0.039000 | There's a bus stop ***in front of the [School name] school.*** |
| **14** | 0.038636 | Humanities at ***[Place name]*** and [Place name] Campus |
| **15** | 0.038547 | That gender something ***[Person's name]*** professor ***at [University name]*** |
| **16** | 0.038491 | ***The [Railway name]*** line extends about east-northeast. |
| **17** | 0.038466 | I wouldn't mind if we did river trips or something along ***places like the [Place name] River***, but... |
| **18** | 0.038237 | From ***the Jomon*** period to today's Japan |
| **19** | 0.038152 | If it's one-third, I'll teach it like ***three equal parts***. |
| **20** | 0.038132 | ***One litre carton of milk*** |

**Supplementary Table 4. Production PC2 lowest correlation utterances (Self-mentalizing).**

| Rank | Correlation coefficient | Utterance |
|---|---|---|
| **1** | -0.057054 | ***It's like, yes, I know.*** |
| **2** | -0.056760 | ***I think it's going to be a lot less*** strange. |
| **3** | -0.056570 | ***it may be.*** |
| **4** | -0.054860 | ***I don't know.*** |
| **5** | -0.054393 | ***I think it was different.*** |
| **6** | -0.053113 | ***I think there are*** situations like that. |
| **7** | -0.053033 | ***I think*** the city ***would be*** delineated. |
| **8** | -0.052673 | That ***may well be the case.*** |
| **9** | -0.052557 | ***I think it's interesting.*** |
| **10** | -0.052359 | ***Is that so? I don't know.*** |
| **11** | -0.052318 | ***I think*** he ***is*** a first-class person. |
| **12** | -0.052026 | ***I guess I can*** trust you a little on that**.** |
| **13** | -0.051964 | ***I think it's just that*** they are too strict. |
| **14** | -0.051216 | ***I'm not*** that worried**.** |
| **15** | -0.050767 | ***I think people might think*** it's a time of hesitation. |
| **16** | -0.050167 | ***I think it would be*** quite ***difficult to*** go to the spa. |
| **17** | -0.049680 | ***I think it's totally different.*** |
| **18** | -0.049657 | ***I think they'll do whatever it takes.*** |
| **19** | -0.049388 | ***It sounds like you've been*** going there ***recently***. |
| **20** | -0.049295 | ***I think I'm going to*** get married. |

**Supplementary Table 5. Production PC3 highest correlation utterances (Sentence construction).**

| Rank | Correlation coefficient | Utterance |
|---|---|---|
| **1** | 0.030379 | ***Yes, on the same side*** it is. |
| **2** | 0.030345 | If you are told to do this, ***you basically say yes*** and do it. |
| **3** | 0.029521 | I don't know how to get there ***from the front gate***. |
| **4** | 0.029000 | I think we should ***be able to create a*** spiritual ***base***. |
| **5** | 0.028370 | ***Someone*** from the same department ***was next to me, so*** I chatted with him. |
| **6** | 0.028179 | ***Once*** I say ***yes***, it's often like, "Well, it's not so hard after thinking about it." |
| **7** | 0.027844 | when all animal elements ***are classified*** |
| **8** | 0.027797 | Don't really like ***anything that isn't checked***? |
| **9** | 0.027447 | ***Turn right and*** you'll see ***a parking lot***. |
| **10** | 0.027205 | It's a place called [Place name]. Yes. ***There, yes.*** |
| **11** | 0.026958 | ***It's running as*** a regular use by the general public. |
| **12** | 0.026624 | As ***you approach*** the [Place name] ***Campus***, you will first see Lawson on your left. |
| **13** | 0.026580 | ***Yes, first semester, second semester.*** I'm just a little past the middle of |
| **14** | 0.026517 | ***There is a train coming***, bound for [Train station], so I will take it. |
| **15** | 0.026386 | ***Or the*** creative ***part reacts.*** |
| **16** | 0.026216 | ***I went on to the department*** and became a sophomore |
| **17** | 0.026017 | ***If you go along the river*** on the right side of the road, ***you can get there*** |
| **18** | 0.026013 | I think ***as long as they have a job*** at the very least, that's all that matters. |
| **19** | 0.025933 | I don't take long baths, but ***I stay in until*** I'm properly warm. |
| **20** | 0.025880 | but ***they close the entrance*** for a while or something like that. |

**Supplementary Table 6. Production PC3 lowest correlation utterances (Fillers).**

| Rank | Correlation coefficient | Utterance |
|---|---|---|
| **1** | -0.082116 | ***Uh, hmm*** |
| **2** | -0.081504 | ***Ahhhhhhhh, that one.*** |
| **3** | -0.080672 | ***Uh-oh.*** |
| **4** | -0.075893 | ***hmmmmmm*** |
| **5** | -0.074875 | ***whoo-whoo-hoo!*** |
| **6** | -0.074194 | ***uh-huh*** |
| **7** | -0.073668 | ***Uh ehh...*** |
| **8** | -0.073069 | ***hahahahahahaha!*** |
| **9** | -0.072478 | ***hahahahahaha*** |
| **10** | -0.072028 | ***Hmmm ah.*** |
| **11** | -0.071188 | ***Uh, well...*** |
| **12** | -0.071096 | ***Oh, well, well, well, well.*** |
| **13** | -0.070474 | ***oh what a mess*** |
| **14** | -0.070037 | ***Well, ahm...*** |
| **15** | -0.069135 | ***hahahahaha!*** |
| **16** | -0.069050 | ***a-ha-ha-ha-ha!*** |
| **17** | -0.068962 | ***Uh*** |
| **18** | -0.068774 | ***Eh, hmm.*** |
| **19** | -0.068439 | ***ahaha yeah yeah*** |
| **20** | -0.068160 | ***Heh, uh-oh.*** |

**Supplementary Table 7. Production PC4 highest correlation utterances (Backchannels).**

| **Rank** | **Correlation coefficient** | **Utterance** |
|---|---|---|
| **1** | 0.103767 | ***Yes, yes, yes.*** |
| **2** | 0.103227 | ***Oh, yes, yes, yes.*** |
| **3** | 0.097924 | ***Yes, yes*** |
| **4** | 0.097567 | ***Oh, yes, yes.*** |
| **5** | 0.094828 | ***Yes, right.*** |
| **6** | 0.090814 | ***Yes, sorry*** |
| **7** | 0.090491 | ***Uh, yes.*** |
| **8** | 0.090457 | ***Oh, yes, yes, yes.*** |
| **9** | 0.090327 | ***right, yes.*** |
| **10** | 0.088577 | ***Yes.*** |
| **11** | 0.088467 | ***Yeah.*** |
| **12** | 0.088181 | ***Ah, yes.*** |
| **13** | 0.088099 | ***Yes*** |
| **14** | 0.087895 | ***Uh, yes, that's right.*** |
| **15** | 0.087405 | ***Is it? Yes*** |
| **16** | 0.087365 | ***Well, yes.*** |
| **17** | 0.087006 | ***Yes, there is.*** |
| **18** | 0.086603 | ***Are there any*** benefits? ***Yeah*** |
| **19** | 0.086428 | ***Yes, excuse me.*** |
| **20** | 0.086355 | ***Yes, yes, I remember.*** |

**Supplementary Table 8. Production PC4 lowest correlation utterances (Ambiguous expressions).**

| Rank | Correlation coefficient | Utterance |
|---|---|---|
| **1** | -0.007837 | ***I have a feeling that I want to try it.*** |
| **2** | -0.005409 | ***Reading*** to gain knowledg**e *is somehow*** different. |
| **3** | -0.004374 | I was thinking of ***doing*** various things ***and having fun.*** |
| **4** | -0.003635 | ***At one time,*** that kind of thing was ***a really weird thing to say.*** |
| **5** | -0.003462 | ***So it's a little something.*** |
| **6** | -0.003259 | ***They were having something like that.*** |
| **7** | -0.003220 | ***That's something like...*** |
| **8** | -0.002674 | ***I'm like the person who's been*** criticizing people so far. |
| **9** | -0.002630 | ***It's easier*** to move around that way. |
| **10** | -0.002396 | ***I think the best thing to do is*** to get the vaccine**.** |
| **11** | -0.001671 | I've usually been able to jump on ***something like that.*** |
| **12** | -0.001651 | A friend told me there was ***something like this.*** |
| **13** | -0.001650 | I'm sorry I didn't mean it in the way I said it, ***that he's just trying to*** get a laugh out of them. |
| **14** | -0.001631 | ***It's like a weird feeling in my stomach.*** |
| **15** | -0.001451 | ***I don't like that kind of thing.*** |
| **16** | -0.001287 | ***On the contrary,*** I can't stop because it's kind of fun. |
| **17** | -0.001197 | ***in that sense*** |
| **18** | -0.001140 | ***Some kind of*** intermediate between materials and light or ***something like that.*** |
| **19** | -0.001019 | ***But*** there's nothing to do if I stay over there ***that long.*** |
| **20** | -0.000853 | If ***something like that*** goes away. |

**Supplementary Table 9. Comprehension PC1 highest correlation utterances (Turn-taking).**

| Rank | Correlation coefficient | Utterance |
|---|---|---|
| **1** | 0.169372 | I'm sure ***they*** probably ***do*** a lot of thinking about that, too. |
| **2** | 0.169140 | There is a point where ***I wonder if I should*** treat human beings in a special way. |
| **3** | 0.169045 | I could tell right away that ***it was the right one****.* |
| **4** | 0.168803 | I'm talking about a TV show called [TV program name] |
| **5** | 0.168778 | This is a program ***that*** tries to clarify why this is a bus route that only runs once a year. |
| **6** | 0.168675 | I think it's ***something that*** should be avoided. |
| **7** | 0.167813 | ***And*** to see if we can see ***the*** patterns properly or if they change. |
| **8** | 0.167516 | I often read really useless books that my parents ***would tell me not to do*** that if they saw them. |
| **9** | 0.167406 | ***I don't*** recommend it, ***though***. |
| **10** | 0.167355 | These books ***often*** deal with what makes them different from others. |
| **11** | 0.167318 | I'm talking ***about*** how we pack when we shop. |
| **12** | 0.167216 | I've heard that a person's judgment and a lie detector's judgment ***aren't that different****.* |
| **13** | 0.167056 | People who come to consult with the doctor ***about their inability*** to stop eating. |
| **14** | 0.167009 | ***I haven't even had*** a barbecue recently anymore. |
| **15** | 0.166743 | I play a lot of games, ***but*** when I do that... |
| **16** | 0.166686 | The inspections seem a little close, ***but*** there are some places that are ***a little far away****.* |
| **17** | 0.166587 | ***Let's*** think about the reason. ***It's*** just in ***this story****.* |
| **18** | 0.166574 | ***I don't care if*** she’s a real mom, ***but*** if ***you're talking about doing it****.* |
| **19** | 0.166515 | ***First of all,*** I would like to ask you how ***you would like to put it in.*** |
| **20** | 0.166394 | What is it ***that separates*** chimpanzees and people? |

**Supplementary Table 10. Comprehension PC1 lowest correlation utterances (Turn-taking).**

| Rank | Correlation coefficient | Utterance |
|---|---|---|
| **1** | 0.104871 | ***Shu, Shu**,* what kanji, Shuren |
| **2** | 0.106646 | ***Oh, yeah.*** |
| **3** | 0.107250 | To ***[Railway name]*** in [Train station name]. |
| **4** | 0.107430 | ***Hahaha, yeah.*** |
| **5** | 0.107508 | Monorail, that line, is not ***east*.** |
| **6** | 0.107635 | It's ***easier*** to decide from those that seem to have the highest priority. |
| **7** | 0.107759 | I'm doing it now in a more senior tone, as a pattern that makes it harder to ***refuse*.** |
| **8** | 0.107876 | Is it north of [Train station], or ***north*** of the monorail? South? |
| **9** | 0.107922 | I feel like I'm just now learning about the ***harsh*** world of math. |
| **10** | 0.107998 | ***Against*** drunk driving, I think it's not good. |
| **11** | 0.108046 | ***West*** |
| **12** | 0.108079 | ***Haha, yeah.*** |
| **13** | 0.108081 | Ishikawa is pretty much like ***a plain.*** |
| **14** | 0.108094 | When you want to say that you are not available in a subtle way. ***Ah*, *or*** maybe |
| **15** | 0.108258 | What was it, ***the ocean,*** what happens when the sea level rises. |
| **16** | 0.108340 | ***crane*** and tortoise |
| **17** | 0.108474 | ***Sea, yeah*** |
| **18** | 0.108503 | ***Yeah, yeah, right.*** |
| **19** | 0.108693 | ***Eighteen*** |
| **20** | 0.108697 | ***Pacific Ocean*** |

**Supplementary Table 11. Comprehension PC2 highest correlation utterances (Backchannels or involuntary laughter).**

| Rank | Correlation coefficient | Utterance [Original Japanese] |
|---|---|---|
| **1** | 0.141284 | ***Uh-huh-muh-muh [ah un un un un]*** |
| **2** | 0.140221 | ***hehehehehehehe [fu fu fu fu fu fu fu]*** |
| **3** | 0.137295 | ***ahahahahahahahaha*** |
| **4** | 0.137209 | ***Oh, yeah, yeah, yeah, yeah, yeah. [oh un un un un un]*** |
| **5** | 0.136622 | ***ahahahahaha*** |
| **6** | 0.136158 | ***Uh, yeah, yeah, yeah. [ah un un un]*** |
| **7** | 0.135765 | ***Ahahahahahahahaha*** |
| **8** | 0.135190 | ***Oh, yeah, yeah, yeah. [ah un un un]*** |
| **9** | 0.133566 | ***hehehehehe [fu fu fu fu fu]*** |
| **10** | 0.133247 | ***ahahahahahaha*** |
| **11** | 0.130119 | ***ha ha ha ha yeah [ahahaha un]*** |
| **12** | 0.129987 | ***ahahahahaha*** |
| **13** | 0.129437 | ***hahahahahaha*** |
| **14** | 0.128591 | ***Hahahahahahaha*** |
| **15** | 0.127973 | ***Ahahahahaha*** |
| **16** | 0.127358 | ***hehehehe [fu fu fu fu]*** |
| **17** | 0.126450 | ***uh-huh-huh [ah un uhn]*** |
| **18** | 0.126280 | ***hahahahaha*** |
| **19** | 0.125718 | ***Uh huh-huh [ah un un]*** |
| **20** | 0.125697 | ***Uh-huh-huh-huh [ah un un un]*** |

**Supplementary Table 12. Comprehension PC2 lowest correlation utterances (High-syllable content words).**

| Rank | Correlation coefficient | Utterance |
|---|---|---|
| **1** | 0.005959 | And if not, it's going to mean a job in a corporate position, ***like a college position***. |
| **2** | 0.006466 | ***For example,*** when ***you make this into a study or something.*** |
| **3** | 0.009715 | ***Our schools*** and jobs, ***for example***, are usually a one-year unit. |
| **4** | 0.010032 | ***I graduated from the [School name]*** |
| **5** | 0.011129 | Everyone gets ***your face*** now, ***for example.*** |
| **6** | 0.011350 | Maybe a teacher at the university, or ***a club member, or someone at a part-time job***. |
| **7** | 0.011615 | ***When I worked for my*** last ***company***. |
| **8** | 0.012085 | If it were you, you might ***read books on philosophy*** or something like that. |
| **9** | 0.012176 | What do you do ***when recruiting*** locally, for example? |
| **10** | 0.012359 | I wanted to be in ***the clothing industry, or rather,*** I wanted to be in ***the fashion industry.*** |
| **11** | 0.012687 | There are many people who are working hard in academia ***or in the corporate world.*** |
| **12** | 0.012696 | ***The kind of*** thing ***they teach in college*** is already in the books. |
| **13** | 0.012749 | I think ***having corporate experience*** is not a bad thing. |
| **14** | 0.013028 | What about ***people in*** other ***faculties*** or departments? |
| **15** | 0.013601 | It seems that there was ***a meeting*** to be held first thing in the afternoon |
| **16** | 0.013901 | Maybe you can imagine what it would be like ***if you went into*** a private tutor or something. |
| **17** | 0.014057 | ***The image*** of such a person is almost always connected to the image of ***the place***. |
| **18** | 0.014099 | Maybe ***a teacher at the university, or*** a club member, or someone at a part-time job. |
| **19** | 0.014327 | Like where you are now in relation to ***your future self or something.*** |
| **20** | 0.014434 | ***The book for undergraduates*** was surprisingly interesting. |

**Supplementary Table 13. Comprehension PC3 highest correlation utterances (Utterance-initial fillers or words).**

| Rank | Correlation coefficient | Utterance |
|---|---|---|
| **1** | 0.069043 | Uh, ***well,*** I have a favour to ask you, could you please… |
| **2** | 0.068731 | (Start of the run) ***Well***, I would like to ask you about the way you say things… |
| **3** | 0.065117 | (Start of the run) ***Well***, I was wondering if you could tell me what the students think about the classes. |
| **4** | 0.065083 | (Start of the run) ***Let's see***, do you remember the story of Cinderella? |
| **5** | 0.064332 | (Participant’s turn: “Real daughter and daughter-in-law, uhhhh…”**)** Oh, I see. ***Uh, well,*** in Cinderella's parents… |
| **6** | 0.063679 | (Start of the run) ***Say, um,*** well, I was told it's time for tests, but there are hardly any reports or anything like that? |
| **7** | 0.061238 | (Start the topic in the beginng part of the run) ***Well, let's see...*** about ***this upcoming*** theme... |
| **8** | 0.061192 | (Start of the run) ***Let's see,*** this is the first one that appeared in a logic book or something. |
| **9** | 0.060605 | (Start of the run) ***Let’s see,*** I think we did arithmetic addition in elementary school. |
| **10** | 0.058321 | (Start of the run) ***Okay, so first,*** self-introductions, um, which one of us would like to start with? |
| **11** | 0.058164 | (Start of the run) ***Let’s see,*** it's a bit like a guessing game. |
| **12** | 0.058158 | (Start of the run) Okay, well, I'd like to start by self-introductions. ***Well...*** could you please begin? |
| **13** | 0.058093 | One more thing...let me ask you about the main theme of this issue***...well***, it may be ***more*** fanciful, |
| **14** | 0.058052 | Ah, yes. I understand. ***Let's see, then,*** could you just list them? |
| **15** | 0.057820 | (Start of the run) ***Okay, well,*** let me start with a question. |
| **16** | 0.057597 | (Start of the run) ***Well, uh, then,*** could you please broach the subject? |
| **17** | 0.057361 | (Start of the run) ***Okay, so*** here's the situation we’re in the jungle in… |
| **18** | 0.057260 | we are influenced by our surroundings, ***so well, you know,*** it is easier to achieve goals when you… |
| **19** | 0.057179 | (Start of the run) Okay, ***so, well,*** I was just asking you, is it a book or something? |
| **20** | 0.057081 | (Start of the run) I’m sorry, but I explained that we hadn't started scanning yet. ***Well, then*** could you please start? |

**Supplementary Table 14. Comprehension PC3 lowest correlation utterances (Sentence construction).**

| Rank | Correlation coefficient | Utterance |
|---|---|---|
| **1** | -0.037736 | No, ***I don't think it's changed.*** |
| **2** | -0.036200 | ***You never know*** *what* **might happen.** |
| **3** | -0.036142 | ***I guess they feel like*** they're not going to miss out. |
| **4** | -0.033446 | ***I think it's not good.*** |
| **5** | -0.033026 | I was talking, ***feeling like I wasn't*** answering my own question. |
| **6** | -0.032914 | ***I didn't feel uncomfortable when*** I was doing it with apples and oranges. |
| **7** | -0.032504 | ***it would be*** more evolutionarily ***correct*** to be discovered as often as possible |
| **8** | -0.031575 | ***It must have been very difficult*** because there were still a lot of tourists. |
| **9** | -0.031397 | ***Don't we have to eat*** strawberries***?*** |
| **10** | -0.031395 | ***I think*** the amount ***would be more*** if you got it in your retirement plan than if you |
| **11** | -0.031283 | I don't know if I remember it or not***, but I don't think I do.*** |
| **12** | -0.031174 | You think ***it would have been better*** to get out of that kind of house and |
| **13** | -0.031112 | ***You didn't do anything that*** would have made such a watch unusable. |
| **14** | -0.031080 | ***I feel like I'm obligated*** to eat. |
| **15** | -0.030726 | ***You might get different*** customer service with someone who looks poor and |
| **16** | -0.030606 | I thought ***it wouldn't matter*** if it wasn't that day. |
| **17** | -0.030425 | I'm sure ***they will be offended by the discrimination.*** |
| **18** | -0.030110 | ***I don't think I've been on it*** for about 30 years. |
| **19** | -0.030002 | I think ***they could have*** imitated it. |
| **20** | -0.029951 | I thought ***it wasn't what*** a researcher ***would say***. |

**Supplementary Table 15. Comprehension PC4 highest correlation utterances (Adverbial phrases).**

| Rank | Correlation coefficient | Utterance |
|---|---|---|
| **1** | 0.041403 | ***Traditionally*** in [Place name]. |
| **2** | 0.037374 | It's a multi-year plan, ***originally***. |
| **3** | 0.036015 | Students are, ***well, more*** sensitive. |
| **4** | 0.035805 | They see you as ***lazy***. |
| **5** | 0.035640 | I think it's on a decade basis, ***essentially*.** |
| **6** | 0.035398 | And if there's a possibility of doing some programming or something ***to some extent*.** |
| **7** | 0.035254 | It's going in an uninteresting direction, ***as a result.*** |
| **8** | 0.035187 | ***Until then*** |
| **9** | 0.035143 | Then ***you don't have to live in [Place name]***, you can live somewhere else at least once. |
| **10** | 0.035053 | You know, this was ***originally*** a book. |
| **11** | 0.034999 | ***Now*** if there was a system |
| **12** | 0.034987 | ***Now that you know*** what you're talking about... |
| **13** | 0.034963 | There's ***a coastline*** and then it goes into the mountains. |
| **14** | 0.034884 | ***Ah, just****,* but I'm not so sure. |
| **15** | 0.034669 | ***Oh, I knew it.*** |
| **16** | 0.034391 | Oh, it's the same as this one, ***earlier, hmmm****.* |
| **17** | 0.034333 | Well, ***depending on*** the university |
| **18** | 0.034210 | Comparatively, ***though, in that*** |
| **19** | 0.034158 | Which was it ***until then***? |
| **20** | 0.034133 | you don't do it ***originally*.** |

**Supplementary Table 16. Comprehension PC4 lowest correlation utterances (Self-mentalizing).**

| Rank | Correlation coefficient | Utterance |
|---|---|---|
| 1 | -0.059888 | ***I just wanted to ask you*** what ***you think.*** |
| 2 | -0.054070 | ***Could you*** start***, please? Please do.*** |
| 3 | -0.053054 | ***Do you understand*** the causal relationship***?*** |
| 4 | -0.052568 | ***Have you heard*** the story***?*** |
| 5 | -0.052563 | ***What do you think is going to happen?*** |
| 6 | -0.052448 | ***Do you have any advice?*** |
| 7 | -0.051729 | ***What do you think?*** |
| 8 | -0.048785 | ***May I ask you something?*** |
| 9 | -0.048068 | ***What do you talk about?*** |
| 10 | -0.048038 | ***I'd like to ask you something.*** |
| 11 | -0.047715 | ***Can you tell me about it?*** |
| 12 | -0.047391 | ***I'd like to ask you something.*** |
| 13 | -0.047238 | ***Did I ask you about it?*** |
| 14 | -0.046892 | ***I'd like to ask you something.*** |
| 15 | -0.046530 | ***I'd like to ask you.*** |
| 16 | -0.046254 | ***I need to ask you something.*** |
| 17 | -0.046110 | ***Have you heard of it?*** |
| 18 | -0.045595 | ***I'd like to ask you something.*** |
| 19 | -0.044977 | ***I'd like to ask you something.*** |
| 20 | -0.044397 | ***Can I ask you about...*** |

**Supplementary Table 17. Comprehension PC5 highest correlation utterances (Concrete nouns).**

| Rank | Correlation coefficient | Utterance |
|---|---|---|
| **1** | 0.061080 | ***[Place name] at [Place name]*** |
| **2** | 0.060908 | ***a litre of milk*** |
| **3** | 0.059717 | ***Apples and oranges*** |
| **4** | 0.059624 | ***and two oranges*** |
| **5** | 0.059581 | ***two potatoes*** |
| **6** | 0.059296 | ***Yes, yes*** |
| **7** | 0.058737 | ***Carrots and rabbits.*** |
| **8** | 0.058652 | ***A pack of milk is*** |
| **9** | 0.058308 | ***Ten eggs.*** |
| **10** | 0.058108 | ***three apples*** |
| **11** | 0.057796 | ***Three apples.*** |
| **12** | 0.057609 | ***Apples and oranges separately*** |
| **13** | 0.057550 | ***Ten eggs in a pack.*** |
| **14** | 0.057489 | ***Two potatoes*** |
| **15** | 0.057282 | ***apples and oranges*** |
| **16** | 0.057171 | ***apples and oranges*** |
| **17** | 0.057117 | ***Crane game, yeah.*** |
| **18** | 0.057038 | ***A bag of potato chips*** |
| **19** | 0.056998 | ***Part-time lecturer*** |
| **20** | 0.056903 | ***bus, yeah*** |

**Supplementary Table 18. Comprehension PC5 lowest correlation utterances (Non-lexical sounds/voluntary laughter).**

| Rank | Correlation coefficient | Utterance [Original Japanese] |
|---|---|---|
| **1** | -0.027070 | ***Ahh ahh ahh ahh*** |
| **2** | -0.025386 | ***Ahahahahaha*** |
| **3** | -0.024444 | ***Ahh ahh ahh*** |
| **4** | -0.022197 | ***hahahahaha [he he he he he]*** |
| **5** | -0.020334 | ***Oh, I've done it!*** |
| **6** | -0.020160 | ***hahahahaha*** |
| **7** | -0.020128 | ***Ahh ahh ah*** |
| **8** | -0.019206 | ***hahahahahahaha [fu fu fu fu fu fu fu fu]*** |
| **9** | -0.019150 | ***Ahahahahahaha.*** |
| **10** | -0.017677 | ***Ahh ahh*** |
| **11** | -0.017529 | ***hahahahaha*** |
| **12** | -0.017159 | ***Hahahahahaha*** |
| **13** | -0.017052 | ***Ahahahahaha*** |
| **14** | -0.016726 | ***Ahahahahaha*** |
| **15** | -0.016558 | ***Woohahahaha*** |
| **16** | -0.015752 | ***Ah, well, I see*** |
| **17** | -0.015377 | ***hahahahaha*** |
| **18** | -0.015202 | ***Oh, I see.*** |
| **19** | -0.014522 | ***Eh, uh, then.*** |
| **20** | -0.014155 | ***Oh, well then...*** |

**Supplementary Table 19. Comprehension PC6 highest correlation utterances (Something the listener may not know).**

| Rank | Correlation coefficient | Utterance |
|---|---|---|
| **1** | 0.048392 | I've heard that the decision-making ability ***doesn't change that much****.* |
| **2** | 0.048320 | ***I'm just wondering*** why it's a bus route that only runs once a year. |
| **3** | 0.048130 | ***what's*** different from the rest of them |
| **4** | 0.047088 | How is it that it just ***pops up****?* |
| **5** | 0.046535 | I wanted to ask you how ***you think about it.*** |
| **6** | 0.046219 | ***It's like saying***, "If you can avoid it, maybe you should avoid it." |
| **7** | 0.045835 | ***I was just wondering*** how ***you think about it.*** |
| **8** | 0.045800 | ***To be seen*** singing*...* |
| **9** | 0.045502 | This game allows ***you to see*** what you want to keep and what you don't want to keep. |
| **10** | 0.045279 | My research theme was to find out why we couldn't do that and how ***we could do it.*** |
| **11** | 0.044140 | ***It*** could have ***been*** divided into groups. |
| **12** | 0.043850 | If the fever rises, it sounds like ***it could be*** a cold. |
| **13** | 0.043203 | ***I'm just wondering*** what it's like. |
| **14** | 0.042901 | ***I was thinking that*** we would like to think about what it would be like. |
| **15** | 0.042755 | The relationship is ***not so*** far apart ***that*** it feels like a rivalry. |
| **16** | 0.042733 | The advantage of online is ***that you can do it*** remotely. |
| **17** | 0.042698 | It's not ***that there are things*** I regret and decide not to do anymore. |
| **18** | 0.042451 | ***I'm just wondering if*** a distinction needs to be made. |
| **19** | 0.042283 | It's difficult, isn't it, ***trying*** not to move it? |
| **20** | 0.041508 | Do you even have a person ***that you would like to be*** like? |

**Supplementary Table 20. Comprehension PC6 lowest correlation utterances (Number/calculation).**

| Rank | Correlation coefficient | Utterance |
|---|---|---|
| **1** | -0.045627 | That would be ***one third divided by two.*** |
| **2** | -0.045037 | That would mean ***there are two thirds.*** |
| **3** | -0.044400 | ***Autumn or Spring Equinoxes*** |
| **4** | -0.041409 | ***Six hundred thousand and one yen*** |
| **5** | -0.041153 | Further halve what is ***still one-third*** done |
| **6** | -0.040975 | ***One half multiplied by*** one third |
| **7** | -0.040647 | ***Six hundred thousand and one thousand yen*** |
| **8** | -0.040646 | One third ***times one half*** |
| **9** | -0.040556 | Multiply ***a third part by*** |
| **10** | -0.040378 | ***One third multiplied by*** one half |
| **11** | -0.040325 | I have now decided on 1, 2, ***3, 4, 5, 5!*** |
| **12** | -0.040288 | ***10 eggs in a package*** |
| **13** | -0.040273 | ***Two potatoes*** |
| **14** | -0.040196 | ***Are you*** moving ***in February or March?*** |
| **15** | -0.039988 | ***One third*** |
| **16** | -0.039657 | ***A half*** |
| **17** | -0.039201 | ***One onion*** |
| **18** | -0.038933 | ***First year in high school*** |
| **19** | -0.038819 | ***One onion*** |
| **20** | -0.038674 | ***Three apples*** |
